# A human-in-the-loop explanation framework for morphologically transparent AI predictions from whole-slide images

**DOI:** 10.64898/2026.01.27.701796

**Authors:** Peiliang Lou, Yitan Zhu, Nicholas Chia, Roopa Kumari, William Yang, Yan Wang, Brenna C. Novotny, Stacey J. Winham, Ruifeng Guo, Ellen L. Goode, Yajue Huang, Wenchao Han, Tianshu Feng, Chen Wang

## Abstract

Deep learning models enable the prediction of clinical endpoints from whole-slide images (WSIs), but many such models function as “black boxes”, lacking transparency about whether and which histomorphological patterns drive their predictions, hindering interpretability and clinical adoption. Here we propose a human-in-the-loop explanation framework, MorphoXAI, which provides both local and global interpretability for deep learning models by incorporating human-expert interpretations. At the global level, it reveals the histomorphological patterns on which the model consistently relies to distinguish between classes of WSIs, as well as the patterns associated with confusion between classes. At the local level, it indicates which of these patterns are used in the prediction of an individual WSI and which regions within the slide correspond to such patterns. We validated our method on a deep learning model trained for ovarian tumor histologic subtype prediction. The results show that our framework generates explanations that accurately reflect the histomorphology underlying the model’s predictions at both global and local levels. For interpretability and clinical utility in diagnostic contexts, human evaluation results showed that our explanations were easy to interpret, rich in diagnostic features, and directly helpful for diagnostic decision-making, thereby enhancing pathologist-AI collaboration. Our work highlights that unifying global and local explanations and grounding them in expert-interpreted morphology enhances the interpretability and verifiability of deep learning models, thereby facilitating the transparent deployment of such models in clinical practice.

## 1 Introduction

Deep learning has enabled the automated analysis of whole-slide images (WSIs) for predicting clinically relevant endpoints, such as cancer subtype, patient survival, or genetic mutations^1^. Despite the predictive performance, it remains unclear whether and which histomorphological patterns deep learning models rely on when making predictions^2–6^ (Fig. 1a). This lack of transparency makes it difficult for pathologists to understand and verify model predictions in diagnostic contexts, hindering the clinical adoption of such models. In response, recent research in explainable artificial intelligence (XAI)^7^ has sought to identify the histomorphological patterns driving model predictions.

**Figure 1.**
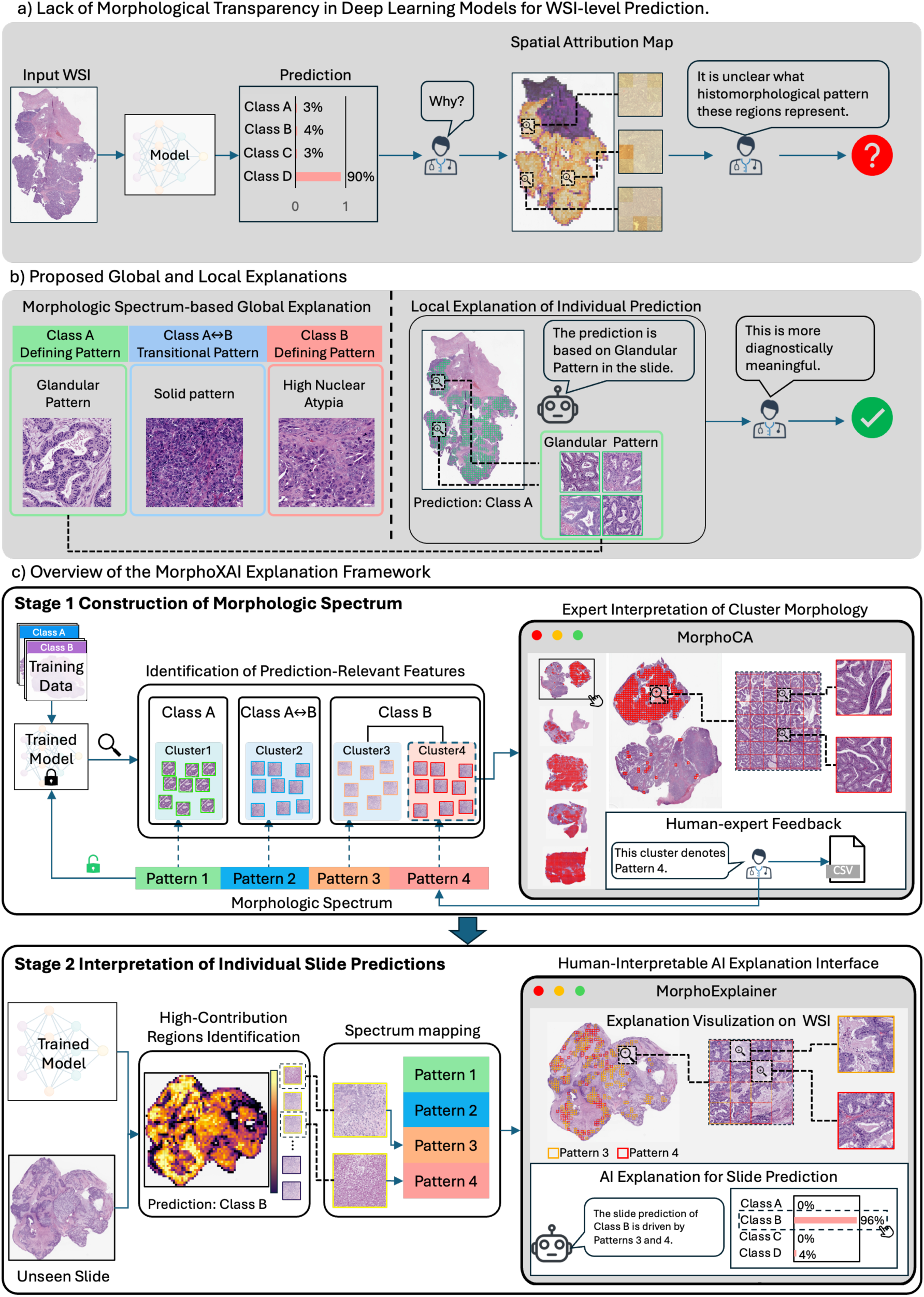
Overview of this work. **a)** Illustration of the lack of morphological transparency in deep learning models for whole-slide image (WSI)–level prediction. **b)** Left: The morphologic spectrum–based global explanation summarizes the key histomorphological patterns the model relies on, including class-defining patterns (e.g., Class A, Class B) and Class A↔B transitional patterns, which correspond to patterns associated with confusion between these two classes (patterns that cause the model to misclassify A as B or B as A). Right: The local explanation of an individual slide prediction maps high-contribution patches to their corresponding spectrum pattern. Patches highlighted with green bounding boxes represent high-contribution regions identified as the Glandular Pattern, which supports the model’s predicted class. **c)** Overview of the MorphoXAI Explanation Framework.

XAI methods for achieving histomorphology-level transparency typically operate in two steps: i) feature attribution^8–14^, which identifies regions within a slide that highly contribute to the model’s prediction; ii) concept mapping^15–17^, which links these high-contribution regions to domain-specific concepts, specifically histomorphological patterns that are defined in diagnostic guidelines and used as evidence in routine diagnosis. Existing XAI methods provide either local explanations^18,19^ or global explanations^9,20–23^. A local explanation explains the model’s prediction for an individual slide by specifying the histomorphological patterns represented by its high-contribution regions. In contrast, a global explanation summarizes the set of histomorphological patterns that the model has learned during training. This is typically achieved by grouping high-contribution regions from multiple slides into clusters based on their feature embeddings and mapping each cluster to a histomorphological pattern.

However, both types of explanations offer only partial insight into the histomorphological basis of model predictions and may even introduce bias. For local explanations, although they make the prediction for an individual slide interpretable, they do not reveal which histomorphological patterns the model consistently relies on to distinguish one class from another across slides. For global explanations, although they characterize the histomorphological patterns that the model has generally learned, they do not indicate which of these patterns are actually used in the prediction of a particular slide or which regions within that slide correspond to these patterns. Furthermore, in both local and global explanations, the mapping from regions to morphological concepts is conducted in the embedding space, for example, by associating region embeddings with concept embeddings derived from text^19^. However, numerous studies have shown that the embedding-based mapping may introduce bias^24–26^, causing mismatches between the mapped concept and the actual morphology of the region. Accurately explaining the morphology requires human expert interpretations, which represents the ground truth.

In this work, we aim (i) to achieve both local and global transparency for deep learning models, clarifying the histomorphological patterns a model generally relies on when distinguishing among classes and identifying the specific patterns it uses when predicting an individual slide, and (ii) to accurately reflect the histomorphology underlying the model’s predictions through expert interpretations. The proposed global and local explanations are illustrated in Fig. 1b. The global explanation is provided in the form of morphologic spectrum, which organizes class-level discriminative features into class-defining and transitional patterns. Class-defining patterns refer to the morphologies that enable accurate class discrimination, whereas transitional patterns correspond to morphologies that lead to confusion between classes, offering insight into both correct and incorrect classifications. The local explanation explains the slide-level prediction by specifying which patterns in the spectrum are represented by the high-contribution regions within the slide. Taken together, the local explanation provides diagnostically meaningful slide-level evidence that allows users to interpret and verify an individual prediction, whereas the global explanation enables users to interpret the model’s predictive behavior at the class level and to assess whether the class-discriminative features it uses are aligned with medical evidence and sufficiently capture the morphologic distinctiveness of that class.

To achieve our goals, we developed a human-in-the-loop explanation framework, MorphoXAI, which comprises two major stages (Fig. 1c). In the first stage, a feature identification procedure is applied to a trained model to identify groups of features learned during training that are associated with different class labels, including those that define each class and those that contribute to inter-class confusion. These feature groups are then mapped back to their corresponding regions in the original WSIs. Subsequently, through a human–AI interaction tool, MorphoCA, experts examine the regions mapped from each feature group across multiple slides to interpret their shared morphological characteristics and determine the corresponding histomorphological pattern for that group, thereby constructing the morphologic spectrum. In the second stage, the constructed spectrum is used to explain the model’s individual predictions. This is achieved by mapping the high-contribution regions from each slide to their corresponding patterns within the spectrum, thereby generating slide-specific local explanations. Together with the prediction results, the local explanations are presented through another human–AI interaction tool, MorphoExplainer, enabling end users to investigate the model’s predictions within the context of WSIs.

We validated our framework using a CLAM^9^ model trained on an ovarian tumor cohort from Mayo Clinic for WSI-based tumor subtyping. In the resulting morphologic spectrum, each pattern’s morphology was well defined, which not only corresponded to a pathologist-recognized histomorphological pattern but also included detailed morphological descriptions. Based on the constructed spectrum, we subsequently generated local explanations for model predictions on an independent dataset. The validation results showed that the model’s prediction behavior on individual slides aligns with how the morphologic spectrum explains both correct and confused classifications at the global level. Pathologists’ evaluations further confirmed that the local explanations accurately reflect the histomorphology of high-contribution patches. To assess interpretability and clinical utility, we applied the explanations in diagnostic scenarios in which pathologists reviewed model predictions together with the accompanying explanations. The evaluation results showed that the MorphoXAI explanations were easy to interpret, enriched with diagnostically relevant features, and directly helpful for diagnostic decision-making. Taken together, these results highlight that unifying global and local explanations and grounding the explanation in expert-interpreted morphology through a human-in-the-loop process could effectively elucidate the histomorphological basis of model predictions, thereby improving the understandability and verifiability of deep learning models. MorphoXAI has the potential to be applied to various models across diverse WSI-based prediction tasks.

## 2 Results

### 2.1 Overview of MorphoXAI

The MorphoXAI framework is designed to elucidate the histomorphological basis of predictions made by deep learning models for WSI classification, providing interpretability at both the global and local levels. Globally, MorphoXAI explains the histomorphological patterns a model has captured from the training slides and the learned associations between these patterns and specific class labels. This level of interpretability clarifies which histomorphological patterns the model consistently relies on to discriminate among classes. Locally, MorphoXAI explains how these pattern–label associations are applied when predicting an individual WSI. This level of interpretability reveals which specific regions within the slide, and which histomorphologic patterns they exhibit, contributed to the model’s prediction.

MorphoXAI constructs the global explanation based on the principle that the pattern–label associations learned by the model during training can be represented as a morphologic spectrum (Fig. 1b). In this spectrum, morphologies that have strong association with a particular class are categorized as class-defining patterns, while those having associations with more than one class are categorized as transitional patterns. Class-defining patterns could exhibit distinct morphological characteristics that are typical of their corresponding class, and their presence tends to lead the model to produce correct classifications. In contrast, transitional patterns could be morphologically ambiguous, exhibiting intermediate characteristics between the defining patterns of the involved classes (e.g., intermediate levels of nuclear atypia), and their presence tends to induce the model’s confusion between these classes. MorphoXAI constructs the local explanation based on the principle that a model’s prediction for an individual slide is driven by the global pattern–label associations. Based on this, MorphoXAI explains the prediction of a slide by mapping its high-contribution regions to the patterns defined in the morphologic spectrum, specifying which pattern is driving this prediction.

The histomorphological patterns learned by the model from the training data are not directly accessible, as they are implicitly encoded in its network. To obtain the morphologic spectrum that globally explains the model, MorphoXAI follows a four-step procedure (Extended Data Fig. 1). In Step 1, MorphoXAI implements a prediction-stability–based strategy. Specifically, multiple models with the same architecture and hyperparameters as the model to be explained are trained on partially overlapping subsets of the training slides, with each model tested on the remaining slides not used for its training. Across these repeated training–testing processes, each slide is tested multiple times by models trained on different subsets of the data. Slides are then stratified based on the prediction stability into two categories: (i) consistently correct: slides that are consistently predicted as their ground-truth class; (ii) highly variable: slides whose predictions frequently alternate between two (or more) classes.

In Step 2, MorphoXAI extracts candidate regions enriched for class-defining or transitional patterns from the stratified slides. Consistently correct slides are assumed to contain class-defining morphologies. Accordingly, regions that are repeatedly identified as high-contribution to the ground-truth prediction across different models are treated as candidate regions enriched for class-defining patterns. In contrast, highly variable slides are assumed to contain transitional morphologies. For these slides, regions that appear as high-contribution in predictions favoring multiple classes are treated as candidate regions enriched for transitional patterns, as their morphology simultaneously supports multiple class assignments. These candidate regions are then clustered into distinct groups in Step 3 based on their feature embeddings, with each group potentially corresponding to a histomorphological pattern.

In Step 4, the regions in each group are mapped back to their corresponding locations in the original WSIs and visualized for pathologists through the MorphoCA interface. With this interactive tool, experts review these regions in their native WSI context across multiple slides and provide slide-level annotations capturing the morphological features consistently observed among the grouped regions within each slide. The corresponding histomorphological pattern for each group is determined through an enrichment analysis of the pathologists’ annotations across slides. To further mitigate subjectivity in expert interpretation, MorphoXAI performs an additional quantitative analysis of cell-type count distributions within each region group. This quantitative context complements human annotations and helps substantiate the resulting pattern definition.

The constructed morphologic spectrum is then used to explain the model’s predictions for individual slides (Fig. 1c, lower panel). For a given slide, MorphoXAI first identifies its high-contribution regions using a feature attribution technique. These regions are then mapped to the corresponding patterns within the morphologic spectrum through an embedding-based concept mapping procedure, yielding the local explanation for that slide. The local explanation, together with the prediction results, is presented to users through the interactive MorphoExplainer interface. This interface enables users to (i) examine, in the full WSI context, whether the actual morphology of each high-contribution region is consistent with the pattern to which it is mapped; (ii) switch between different prediction classes to inspect the class-specific explanations, thereby facilitating interpretation and verification of the model’s decisions.

### 2.2 Study Design for Evaluating MorphoXAI

To assess the utility of MorphoXAI, we conducted an evaluation study applying the framework to explain the CLAM^9^ model, trained using the Mayo Clinic ovarian tumor cohort for the task of ovarian tumor subtype prediction. CLAM is a deep learning model built on the attention-based multiple instance learning (ABMIL) framework, whose variants are widely used for WSI classification^27^. CLAM represents a suitable model for evaluating MorphoXAI, as its architecture and prediction logic align closely with other ABMIL variants, and it has been adopted in a wide range of WSI-based prediction tasks spanning different disease types and clinical endpoints^28–32^. Accordingly, explaining CLAM offers a representative assessment of MorphoXAI’s applicability to a wide range of deep learning models for WSI classification.

The evaluation study included two ovarian tumor cohorts collected at Mayo Clinic: a training cohort and an independent cohort, derived from different time periods and comprising non-overlapping patient groups. Both cohorts contained four histological subtypes—Serous Borderline Tumor (SBT), Clear Cell Carcinoma (CC), Endometrioid Carcinoma (EC), and High-grade Serous Carcinoma (HGSC). The training cohort consisted of 68 patients contributing a total of 602 WSIs, including 21 patients with SBT (134 WSIs), 18 with HGSC (230 WSIs), 11 with CC (101 WSIs), and 18 with EC (137 WSIs). The independent cohort consisted of 31 patients with 221 WSIs, including 11 patients with SBT (77 WSIs), 8 with HGSC (56 WSIs), 4 with CC (27 WSIs), and 8 with EC (61 WSIs). All cases were treatment-naïve, free of mixed histological subtypes, and had subtype confirmation supported by immunohistochemical markers (see Methods for dataset details).

The evaluation process of MorphoXAI proceeded in two stages. First, the CLAM model was trained on the training cohort, after which the MorphoXAI framework was applied to this trained model to construct the morphologic spectrum. The trained model was then used to generate predictions for the independent cohort. Using the fixed morphologic spectrum derived from the training cohort, MorphoXAI subsequently produced local explanations for each individual prediction in the independent cohort. Based on the constructed morphologic spectrum and the resulting local explanations, the evaluation of MorphoXAI focused on two aspects. First, we examined whether the model’s predictions on the independent cohort were consistently grounded in the patterns defined in the spectrum. Second, in a diagnostic setting, we assessed the interpretability and diagnostic helpfulness of the explanations as perceived by pathologists, evaluating their potential to enhance pathologist–AI collaboration.

In the following sections, we outline each step of the spectrum construction and its intermediate outcomes (Sections 2.3–2.5) following the four-step procedure described in Extended Data Fig. 1. Section 2.6 presents the final morphologic spectrum, and Section 2.7 describes how slide-level explanations were generated and visualized using the MorphoExplainer. Section 2.8 reports the evaluation results.

### 2.3 Prediction-Stability-Based Sample Stratification

This step aims to stratify slides into groups enriched for class-defining or transitional morphologies by quantifying their prediction stability across multiple training–testing runs.

First, we trained a CLAM model using CONCH^33^ as the feature encoder on the entire training cohort with empirically determined hyperparameters (see Methods). This full-data model served as the one to be explained by MorphoXAI (Fig. 2a). To quantify prediction stability for sample stratification, we next trained additional models with the same architecture and hyperparameters as the full-data model on partially overlapping patient-level subsets of the training cohort. Specifically, we generated 50 patient-level train/validation/test splits from the training dataset. For each split, a model was trained and evaluated under the same settings, allowing every slide to be evaluated ten times as unseen test data by models trained on distinct subsets of patients (Fig. 2a, 2b; see Methods for details). The prediction stability of each slide was quantified by the fraction of correct predictions among the ten test runs (Acc). Slides were then categorized into three groups: Consistently Correct (Acc ≥ 0.9), Highly Variable (0.1 < Acc < 0.9), and Consistently Incorrect (Acc ≤ 0.1) (Fig. 2b, 2d).

**Figure 2.**
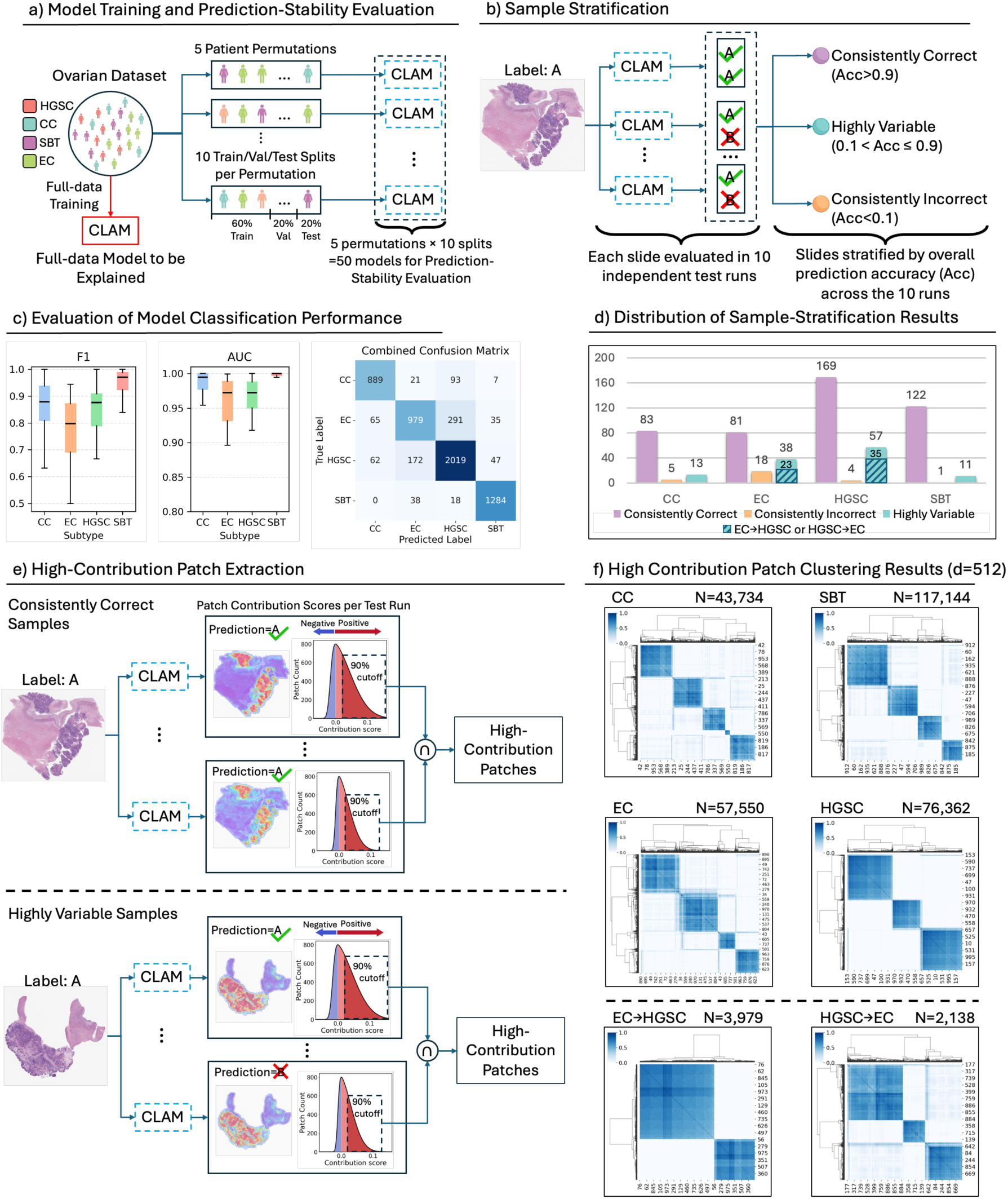
From model training to high-contribution patch clustering: workflow and results. **a)** The model to be explained by MorphoXAI was trained on the entire ovarian dataset (red branch). To estimate slide-level prediction stability, the same dataset was repeatedly partitioned at the patient level to train additional models under identical architecture and hyperparameter settings (blue branch). Specifically, five independent patient permutations were created; for each permutation, patients were partitioned sequentially into ten contiguous blocks, and ten train/validation/test splits (6:2:2) created by cycling the assignment of the ten blocks, so that each split used a different combination of training, validation, and held-out test blocks. Each split trained and evaluated a separate model, yielding 50 independently trained models in total (5 permutations × 10 splits) for prediction-stability evaluation. CC: clear cell carcinoma; SBT: serous borderline tumor; EC: endometrioid carcinoma; HGSC: high-grade serous carcinoma. **b)** Each slide was evaluated ten times as part of the prediction-stability assessment, with each evaluation performed by a model trained on a distinct subset of data. The prediction stability of each slide was quantified by its overall prediction accuracy (Acc)—the fraction of correct predictions among the ten runs—based on which slides were categorized as Consistently Correct, Highly Variable, or Consistently Incorrect. **c)** Boxplots show the distribution of F1 and AUC values from 50 independently trained models. The combined confusion matrix aggregates predictions from ten evaluations per slide, so that the sum of each row equals ten times the number of slides for the corresponding subtype. **d)** Bars show the number of slides in each stability group for each subtype. Within the Highly Variable group, EC→HGSC denotes EC slides frequently misclassified as HGSC, and HGSC→EC denotes HGSC slides frequently misclassified as EC. **e)** Upper panel: method for obtaining high-contribution patches from Consistently Correct slides. Lower panel: method for obtaining high-contribution patches from Highly Variable slides. **f)** Clustering results of high-contribution patches based on embeddings from the CLAM encoder. The top four panels show results for Consistently Correct samples of the four subtypes, while the bottom two panels (EC→HGSC and HGSC→EC) show results for Highly Variable samples whose predictions fluctuated between EC and HGSC. *N* indicates the number of clustered patches; *d* indicates the embedding dimension.

Consistently Correct slides were retained for identifying class-defining patterns across all four subtypes. For Highly Variable slides, most of them originated from cases in which the model confused EC and HGSC (Fig. 2c, 2d). This is consistent with clinical observations, where EC and HGSC are often difficult to distinguish due to the presence of numerous transitional morphologies^34–40^. Accordingly, Highly Variable slides whose predictions alternated exclusively between EC and HGSC were selected for identifying transitional patterns. Slides in the Consistently Incorrect group lacked the tumor-relevant histomorphology required for discovering class-defining or transitional patterns, as they were largely composed of stromal or necrotic tissue with only sparse tumor cells (Supplementary Fig. 1). As such, they were excluded from the analysis.

### 2.4 High-Contribution Patch Extraction and Clustering

This step leverages the attention mechanism of the CLAM model to identify high-contribution regions (patches) within these slides that drive model predictions, enabling localization of class-defining and transitional patterns (Fig. 2e).

The ABMIL model performs slide-level classification by treating each WSI as a bag of smaller image patches (360 μm × 360 μm tiles in this study), each assigned a contribution score quantifying its importance to subtype prediction. Patches with higher contribution scores have a greater influence on the final prediction. Instead of using raw attention weights, MorphoXAI employs a logit-based definition of contribution score, which directly measures the positive or negative impact of each patch on the final classification logits (see Methods for details).

In this ovarian study, high-contribution patches for the subtype prediction were defined as the minimal set of top-ranked patches whose cumulative contributions account for 90% of the total logits for that subtype. Specifically, for the Consistently Correct slides, MorphoXAI identified high-contribution patches associated with the ground-truth subtype across multiple prediction runs and computed their intersection—patches that consistently received high contribution scores in each run and were therefore likely to correspond to class-defining patterns (Fig. 2e, upper panel). For the Highly Variable slides, MorphoXAI extracted high-contribution patches from the prediction sets of both the ground-truth and the confusable subtypes and computed their intersection, capturing regions that alternately supported different subtype predictions and were therefore likely to represent transitional morphologies (Fig. 2e, lower panel).

Next, MorphoXAI grouped patches into clusters using their embedding vectors extracted from the feature encoder of CLAM, which capture morphological and textural characteristics of each patch. Clustering was performed separately for each subtype. To determine the optimal number of clusters within each group, we evaluated multiple candidate numbers *k* ∈ {2, …, 8}, selecting the *k* that achieved maximal clustering stability and consistency (see Methods). The clustering results are summarized in Fig. 2f. For the Consistently Correct samples, we obtained 5 clusters from 43,734 CC high-contribution patches, 4 clusters from 117,144 SBT patches, 4 clusters from 57,550 EC patches, and 3 clusters from 76,362 HGSC patches. For the Highly Variable samples, we obtained 2 clusters from 3,979 patches from EC slides whose predictions fluctuated between EC and HGSC (EC→HGSC), and 3 clusters from 2,138 patches from HGSC slides whose predictions fluctuated between HGSC and EC (HGSC→EC).

### 2.5 Cluster Phenotyping via Human-Expert Interpretation in MorphoCA

This step aims to characterize the histomorphology of each cluster through human-expert interpretations facilitated and collected via the interactive tool MorphoCA.

The workflow is illustrated in Fig. 3a. First, MorphoCA, a custom QuPath extension, streamlines the collection of expert interpretations for cluster phenotyping (Fig. 3b). MorphoCA overlays clustered high-contribution patches on a WSI, allowing experts to switch between clusters and freely zoom in to inspect specific regions while collecting structured annotations for each cluster (see usage instructions in Supplementary Note 1). Considering the potential bias and subjectivity in manual annotations, MorphoXAI further applies CellViT^41^ to quantify cell-type proportions (e.g., tumor, stromal, and necrotic cells) within each cluster, and uses these quantitative profiles to complement pathologists’ annotations, providing additional context that helps substantiate the morphological interpretation of each cluster.

**Figure 3.**
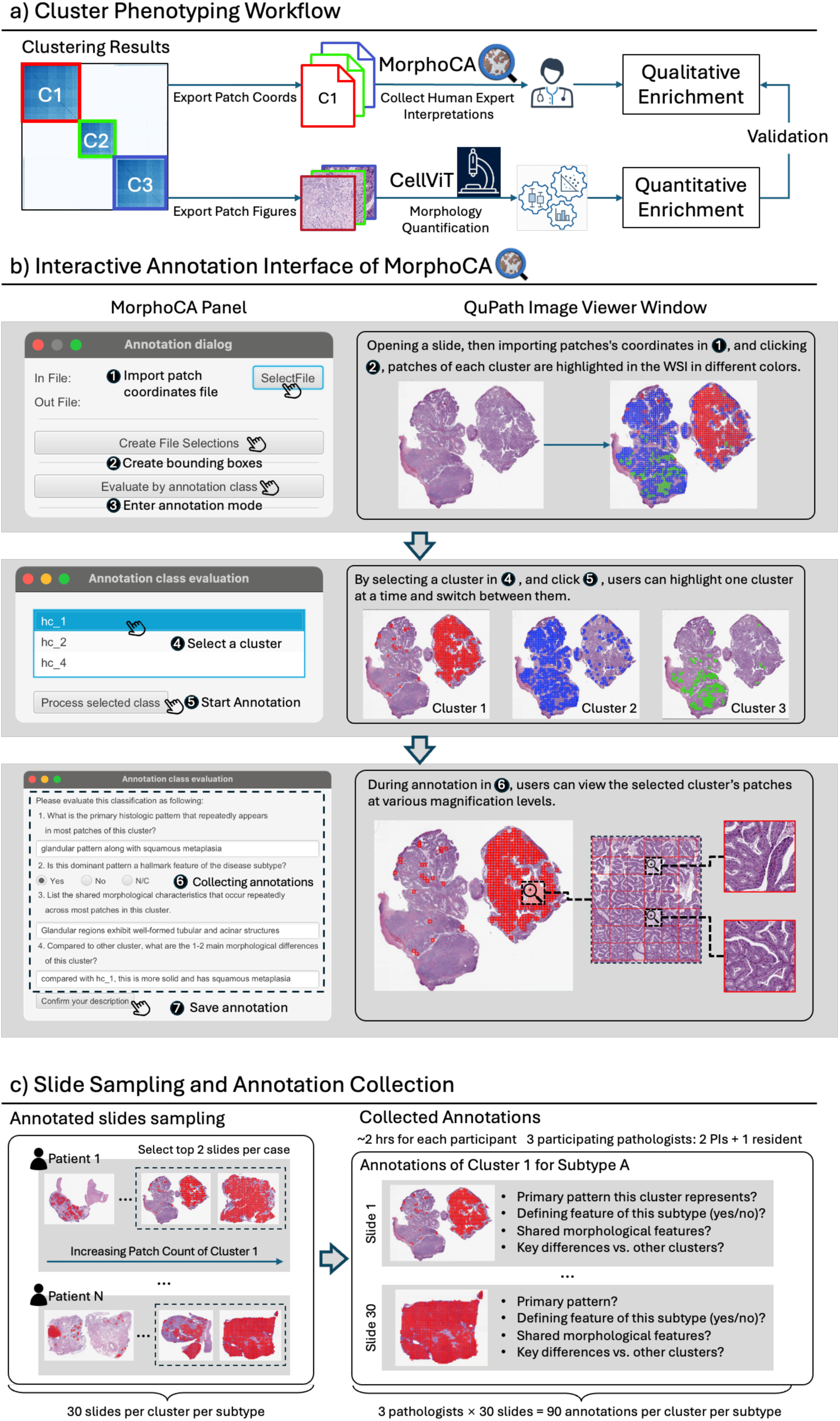
Cluster phenotyping and the collection of human-expert interpretations. **a)** Workflow of cluster phenotyping combining MorphoCA for pathologist annotations with CellViT-based quantitative analysis. **b)** Interactive annotation interface in MorphoCA. Pathologists performed cluster-level annotations within the full-slide context using MorphoCA, a custom QuPath extension designed for interactive visualization and structured annotation. As illustrated, users first open a WSI in QuPath and import a CSV file containing patch coordinates for all clusters. MorphoCA then overlays bounding boxes around patches of each cluster in distinct colors. After entering the annotation mode, users can select one cluster at a time, during which patches from the selected cluster remain highlighted while others are hidden, enabling focused review. A structured annotation panel simultaneously appears, guiding pathologists to record answers to four predefined questions. Annotation results are saved locally upon completion. **c)** For each cluster within a subtype, up to 30 slides were selected for annotation (two slides per patient with the highest number of high-contribution patches; left panel). On each slide, pathologists reviewed the morphology of patches belonging to each cluster and provided structured annotations addressing four predefined aspects: the dominant represented phenotype, whether it constituted a class-defining phenotype, key intra-cluster morphological features, and inter-cluster morphological differences (right panel). Three pathologists (two principal investigators and one resident) participated in the annotation, which yielded 90 annotations per cluster per subtype (3 pathologists × 30 slides), with an average annotation time of approximately 2 hours per pathologist.

In the ovarian study, three pathologists participated in the annotation. To decrease annotation efforts, we adopted the sampling strategy illustrated in Fig. 3c. For each subtype and cluster, slides from each patient were ranked according to the number of high-contribution patches belonging to that cluster. The top two slides per patient were selected, retaining up to 30 slides per cluster for annotation. This sampling ensured that the selected slides captured the majority of high-contribution patches within each cluster while representing morphological diversity across patients. For each selected slide, pathologists sequentially reviewed the patch morphology of every cluster and provided structured annotations addressing four predefined aspects: the dominant represented histomorphological pattern, whether it constituted a subtype-defining pattern, key intra-cluster morphological features, and inter-cluster morphological differences (detailed annotation schema is provided in Methods). In total, this design yielded 90 annotations per cluster (3 pathologists × 30 slides).

### 2.6 Morphologic Spectrum for Explaining Class-Level Predictions of the Ovarian Tumor Subtyping Model

In this section, we present the morphologic spectrum constructed by MorphoXAI, which serves as the global explanation of the CLAM-based ovarian tumor subtyping model. We first introduce the class-defining patterns used to explain the model’s subtype discrimination, followed by the transitional patterns used to explain its confusion between EC and HGSC.

#### 2.6.1 Clear Cell Carcinoma (CC)

MorphoXAI identified five clusters from patches consistently driving the model’s correct CC predictions, and three of them aligned with pathologist-recognized diagnostic patterns, reflecting tubulocystic and hobnail architectures, accompanied by characteristic cytological features such as clear cytoplasm and high nuclear atypia. The other two clusters were associated with necrosis and non-neoplastic tissue, lacking diagnostic relevance for CC. Fig. 4b presents representative tiles from each cluster, visually illustrating their key histomorphological characteristics. Clusters were arbitrarily numbered C1–C5 for identification purposes only.

**Figure 4.**
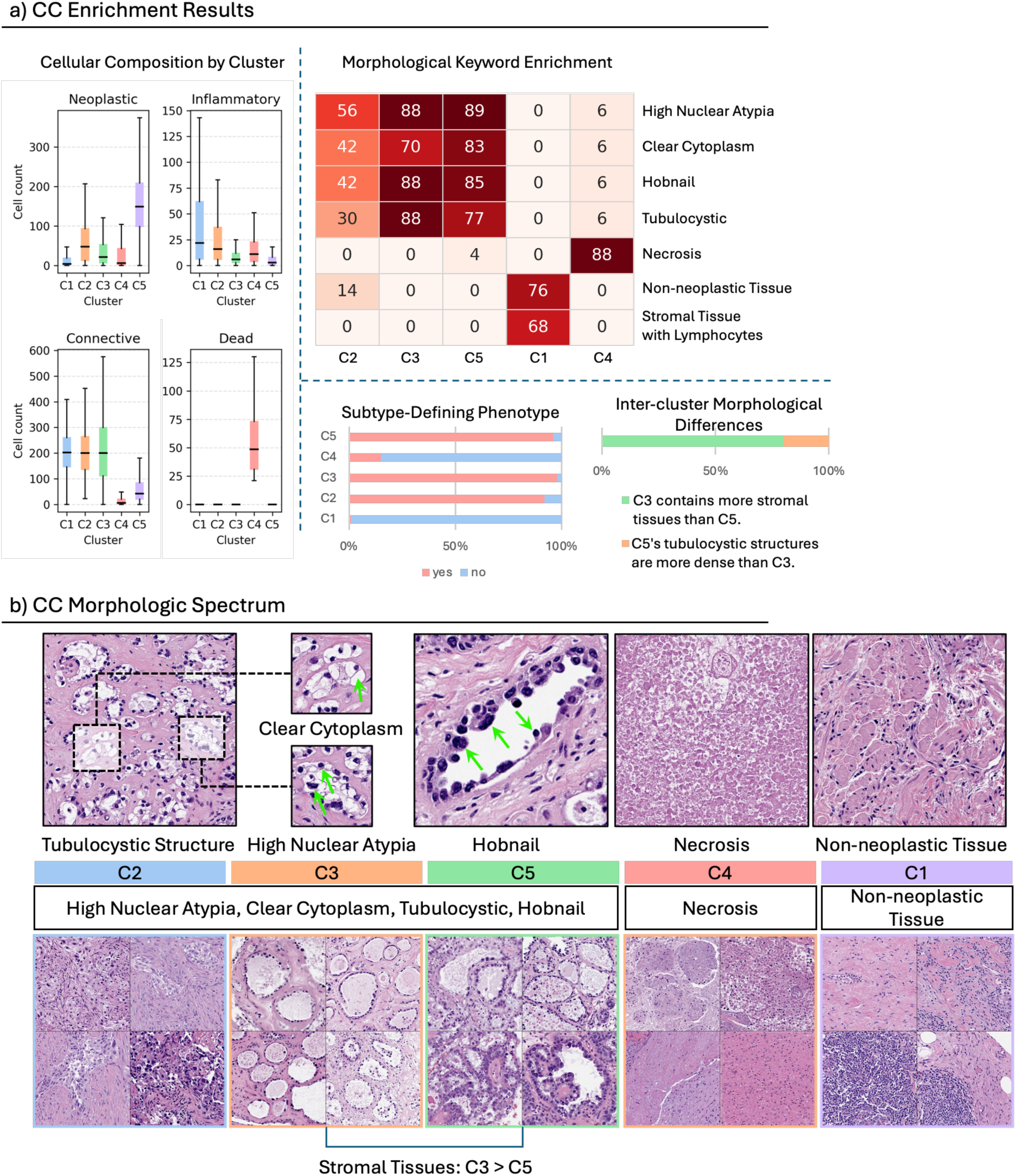
Class-defining patterns for Clear Cell Carcinoma (CC). **a)** Left: Quantitative morphometric analyses assessing the distributions of different cell types within each cluster. Right: Morphological keyword enrichment derived from pathologist annotations for each cluster. The annotations were semantically summarized into keywords, and the resulting heatmap displays keywords as rows and clusters as columns. Each cell’s saliency value represents the number of annotations linking that cluster to the corresponding keyword. Subtype-defining phenotype: Bar plots show the proportion of yes/no responses from pathologists to the question of whether each cluster represents the diagnostic pattern used in clinical practice for CC. Inter-cluster morphological differences: Derived from pathologist annotations describing subtle distinctions among clusters with similar phenotypes. The stacked bar chart shows the percentage distribution of pathologists’ responses for each observed difference. **b)** Representative image galleries illustrating the histomorphological patterns associated with each cluster. Clusters were arbitrarily numbered for identification purposes only. Green arrows highlight specific morphology corresponding to the textual descriptions.

To determine the histomorphology represented by a cluster, MorphoXAI performs enrichment analysis of keywords extracted from pathologist-provided morphological interpretations (Methods). The enrichment results (Fig. 4a, upper right panel) revealed that clusters C2, C3, and C5 were significantly enriched with the keywords “tubulocystic structures,” “hobnail,” “clear cytoplasm,” and “high nuclear atypia.” Visually, these enriched morphological keywords corresponded to two histomorphological patterns (Fig. 4b):

- **Tubulocystic structures**, characterized by glandular or cystic spaces lined by tumor cells. Cells comprising these tubulocystic structures exhibited abundant, glycogen-rich clear cytoplasm, imparting a characteristic clear appearance, and displayed high nuclear atypia, manifested as significant variations in nuclear size, shape, and chromatin patterns.
- **Hobnail structures**, characterized by tumor cells with prominent bulbous nuclei protruding distinctly into glandular lumens, representing a hallmark cellular arrangement frequently observed in clear cell carcinoma.

To clarify subtle distinctions in morphology between clusters that represent the same histomorphological pattern, MorphoXAI collects pathologist feedback on inter-cluster morphological differences. The results (Fig. 4a, lower right panel) indicated the differences between C3 and C5: C3 contained a higher proportion of stromal tissues compared to C5, whereas C5 exhibited denser tubulocystic structures. Cluster C1 was enriched with keywords “non-neoplastic tissue” and “stromal tissue with lymphocytes,” corresponding to predominantly stromal regions lacking diagnostic tumor features. Cluster C4 was specifically associated with “necrosis,” visually represented by extensive necrotic debris and regions devoid of viable tumor cells.

MorphoXAI examines quantitative cell-type distributions within each cluster to corroborate pathologists’ interpretations. The results (Fig. 4a, left panel) showed that Cluster C1 exhibited the lowest tumor-cell percentage, the highest proportion of immune cells, and a relatively high proportion of connective cells, consistent with its enriched keywords, “non-neoplastic tissue” and “stromal tissue with lymphocytes”. C4 showed a significantly higher proportion of necrotic cells compared to other clusters, supporting its unique enrichment of “necrosis.” C2, C3, and C5 displayed high proportions of tumor cells, consistent with their tumor-associated histomorphological patterns (“tubulocystic structures” and “hobnail cells”). Additionally, the quantitative analysis confirmed the subtle morphological differences between C3 and C5, showing a higher proportion of connective cells in C3 and a higher proportion of tumor cells in C5.

To evaluate whether a model-identified class-defining pattern align with diagnostic criteria, MorphoXAI collects pathologist feedback on whether such pattern represents a “Subtype-Defining Phenotype”, defined as a histomorphological pattern specified in diagnostic guidelines and routinely used in clinical practice. The results (Fig. 4a, lower right panel) showed that clusters C2, C3, and C5 are overwhelmingly (>90%) recognized as diagnostic patterns of CC. In contrast, clusters C1 and C4 are predominantly (>75%) recognized as common, non–subtype-specific morphologies that do not provide discriminative diagnostic information for CC. The results (Fig. 4a, lower right panel) show that clusters C2, C3, and C5 are overwhelmingly (>90%) recognized as diagnostic patterns of CC, whereas clusters C1 and C4 are predominantly (>75%) regarded as non-diagnostic, indicating that, globally, the model captures both diagnostic and non-diagnostic morphologies to distinguish CC from other subtypes.

#### 2.6.2 Ovarian High-grade Serous Carcinoma (HGSC)

MorphoXAI identified three clusters from patches consistently driving correct HGSC predictions. Based on the keyword enrichment analysis and the “Subtype-Defining Phenotype” feedback, clusters C2 and C3 corresponded to diagnostic patterns of HGSC (“solid pattern”, “pleomorphism”, “high nuclear-cytoplasmic (N/C) ratio”, “nuclear hyperchromasia”; representative morphology shown in Extended Data Fig. 2b), whereas C1 reflected non-diagnostic stromal regions. Cellular composition analysis further supported these interpretations, revealing high tumor-cell proportions in C2 and C3, and a predominantly stromal composition with minimal tumor cells in C1 (Extended Data Fig. 2a, left panel). Regarding the morphological distinctions between clusters C2 and C3, pathologists did not reach a consistent assessment (Extended Data Fig. 2a, lower right panel). Some judged the two clusters to be morphologically similar, while others noted subtle differences such as more pronounced papillary structures in C3 or fewer tumor cells and increased stromal components in C2. However, these observations were not supported by the quantitative cellular composition analyses; the morphological distinctions between the two clusters remain inconclusive.

#### 2.6.3 Ovarian Serous Borderline Tumor (SBT)

MorphoXAI identified four clusters from patches consistently driving correct SBT predictions (Extended Data Fig. 3b). Clusters C1 and C2 corresponded to diagnostic patterns, each representing a distinct variant of papillary architecture. Specifically, keywords enriched in cluster C1 highlighted papillary structures characterized by multilayered epithelial proliferation and prominent epithelial tufting. In contrast, cluster C2 predominantly represented classic papillary structures lined by a single-layered epithelium with minimal or no cellular stratification or tufting. This morphological distinction was further supported by cellular composition analysis, which revealed a significantly higher proportion of neoplastic cells in cluster C1 compared to C2, consistent with the epithelial proliferation and multilayering observed in C1. C3 and C4 reflected non-diagnostic stromal features (“non-neoplastic tissue,” “ovarian stroma”). Inter-cluster morphological difference annotations highlighted that C3 contained denser stromal tissues compared to C4, a finding quantitatively supported by cellular composition analyses showing a significantly higher proportion of connective cells in C3 than in C4.

#### 2.6.4 Ovarian Endometrioid Carcinoma (EC)

MorphoXAI identified four clusters from patches consistently driving correct EC predictions (Extended Data Fig. 4b). C1, C2, and C4 corresponded to distinct diagnostic patterns: C1 was enriched with squamous metaplasia, C4 predominantly exhibited classical glandular architecture, and C2 showed a mixed diagnostic pattern combining glandular structures with squamous metaplasia. Cellular composition analysis showed consistently high tumor-cell proportions across these diagnostic clusters, and their morphological distinctions were supported by pathologists’ inter-cluster morphological difference annotations. In contrast, C3 was significantly enriched with keywords indicating non-diagnostic patterns (“non-neoplastic tissues,” “stromal tissue,” and “vascular structure”), a finding corroborated by cellular composition analysis showing substantially fewer tumor cells and correspondingly higher proportions of immune and connective cells compared with C1, C2, and C4.

#### 2.6.5 Transitional Patterns Identified from EC→HGSC

This section describes the transitional patterns MorphoXAI identified from EC slides that were frequently misclassified as HGSC (EC→HGSC). Transitional patterns between two subtypes often exhibit histomorphological characteristics that lie between the subtype-defining patterns of each subtype—for example, intermediate degrees of nuclear atypia. To capture these intermediate characteristics, MorphoXAI collects pathologist feedback on the morphological differences between each transitional cluster and the subtype-defining patterns of the corresponding subtypes. In this ovarian study, this evaluation was implemented through the annotation question “Key Morphological Differences from Usual HGSC and EC”. MorphoXAI identified two distinct clusters from the high-contribution patches of the Highly Variable EC→HGSC slides. For each cluster, we collected 69 annotations (3 pathologists × 23 slides).

In summary, clusters C1 and C2 shared certain characteristics with EC and HGSC defining patterns yet also displaying unique morphological distinctions (Extended Data Fig. 5b). Both clusters demonstrated intermediate nuclear atypia, particularly reflected by differences in nuclear size, clearly supporting their transitional status. Regarding the similarities to EC defining patterns: C1 exhibited glandular structures (enriched in “glandular pattern”), whereas C2 displayed solid growth patterns (enriched in “solid growth pattern”) closely resembling squamous metaplasia. However, both clusters also showed distinct differences. Compared with the typical EC glandular pattern, C1 contained glandular regions accompanied by more frequent mitotic figures and apoptotic bodies (enriched in “Mitosis/Apoptotic Bodies”). Moreover, both C1 and C2 displayed more pronounced nuclear atypia, characterized by enlarged nuclei (enriched in “enlarged nuclear size”) and more prominent nucleoli (“prominent nucleoli”), thus appearing morphologically closer to HGSC.

However, compared with the HGSC subtype-defining patterns (HGSC C2 and C3), nuclear atypia in EC→HGSC C1 and C2 was less pronounced, as reflected in the pathologists’ feedback to the “Key Morphological Differences from Usual HGSC and EC” question (Extended Data Fig. 5a, upper right panel). Pathologists reported that nuclei in EC→HGSC C1 and C2 were generally larger and more variable than those in EC but smaller, more rounded, and less pleomorphic than those in HGSC, displaying lower nuclear-to-cytoplasmic ratios and reduced chromatin condensation relative to HGSC. Collectively, these observations indicate that EC→HGSC C1 and C2 exhibit intermediate nuclear morphology between EC and HGSC. Quantitative analyses were further performed to corroborate the intermediate nuclear atypia findings. Specifically, we calculated the area and roundness of tumor cells among three representative clusters: (1) a core HGSC cluster (HGSC C2), characterized by a solid growth pattern and prominent nuclear pleomorphism; (2) a core EC cluster (EC C1), characterized by squamous metaplasia; and (3) the transitional cluster C2 (EC→HGSC). The results (Extended Data Fig. 5a, upper left panel) indicated that, although the nuclear roundness metric alone did not show clear differences among the clusters, nuclear area analysis clearly supported the pathologists’ observations regarding nuclear pleomorphism: the HGSC cluster (C2) exhibited the greatest variability in nuclear size, reflecting pronounced nuclear pleomorphism; the EC cluster (C1) showed the smallest nuclear size variability, consistent with minimal nuclear pleomorphism; and the transitional EC→HGSC cluster (C2) demonstrated intermediate nuclear pleomorphism, with nuclei that were larger and more size-variable than in EC but less variable than in HGSC.

#### 2.6.6 Transitional Patterns Identified from HGSC→EC

MorphoXAI identified three distinct clusters from the high-contribution patches of the Highly Variable HGSC→EC slides (Extended Data Fig. 6b). Notably, transitional patterns identified from HGSC→EC showed substantial morphological similarity with those from EC→HGSC. Specifically: cluster C3 exhibited glandular structures accompanied by fewer mitotic figures and apoptotic bodies, closely mirroring EC→HGSC C1, while cluster C2 predominantly displayed solid growth, highly mimicking EC→HGSC C2. Moreover, both clusters demonstrated intermediate nuclear atypia between typical HGSC and EC, supported by the pathologists’ interpretations and the quantitative morphological analyses. An additional cluster (C1) distinctly differed from these transitional patterns, primarily comprising ovarian stromal tissue (“high stromal content”) intermixed with a small number of typical HGSC tumor cells (“HGSC nuclear features”). This cluster represented regions with limited tumor involvement and did not exhibit morphological features indicative of transitional patterns.

### 2.7 Generating Slide-Level Explanations and Enabling Human Examination via MorphoExplainer

Using the morphologic spectrum established previously, we further generated slide-level explanations for the model’s predictions. We applied this procedure to the independent Mayo Clinic cohort to obtain explanations for all slides in that dataset. Specifically, we applied the full-data model to obtain predictions for each slide and identified the high-contribution patches. These patches were mapped onto the morphologic spectrum using an embedding-based procedure that compared each patch’s feature embedding with the statistical feature distributions of the spectrum clusters. Assignments were made to the closest-matching cluster, while patches lying outside the distributional bounds of all clusters were designated as outliers rather than being forced into an assignment (Methods). This mapping yielded a slide-level explanation that specifies the histomorphological patterns the model relied on for its prediction. The slide-level explanations, together with the model’s predicted probabilities for each subtype, were then presented to human experts through the MorphoExplainer.

MorphoExplainer is a QuPath plugin which allows users to investigate a deep learning model’s slide-level predictions together with the corresponding explanation outputs in the full WSI context (Fig. 5a). It supports visualizing both spectrum-based slide-level explanations and attention heatmaps. The tool overlays either the heatmap or the spectrum-based explanation on the slide, enabling users to switch between the two views and freely zoom in to inspect specific regions. It also supports switching between different class predictions, displaying the corresponding heatmap and spectrum-based explanation for each predicted class, allowing users to examine not only the evidence underlying the model’s top prediction, but also the histomorphologic basis of alternative class predictions. Separate scoring panels are provided to collect users’ feedback. Detailed usage instructions are provided in Supplementary Note 2.

**Figure 5.**
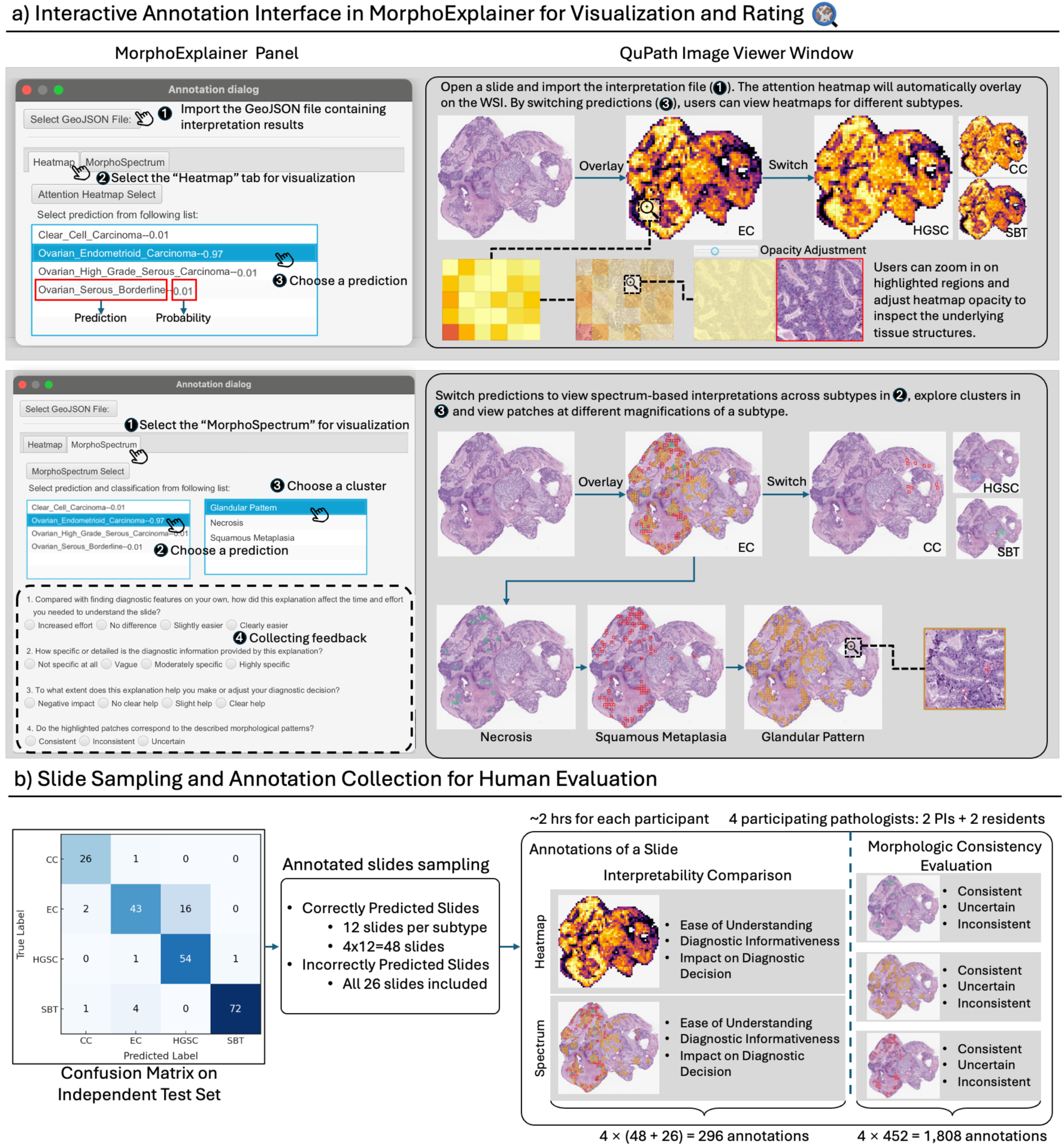
Human evaluation of slide-level explanations using MorphoExplainer. **a)** Interactive annotation interface in MorphoExplainer. Users begin by importing a GeoJSON file containing the model’s predictions, attention heatmaps, and spectrum-based explanations. MorphoExplainer provides two tabs, “Heatmap” and “MorphoSpectrum,” each displaying the predicted classes and their associated probabilities. Pathologists can select any prediction to visualize the corresponding heatmap or spectrum-based interpretation, zoom into regions of interest, and inspect detailed tissue structures. Both tabs include a built-in rating panel that collects user feedback on interpretability, diagnostic specificity, and overall helpfulness for decision-making. **b)** The confusion matrix summarizes the model’s predictions on the independent test set. From these results, 48 correctly predicted slides (12 per subtype) and all 26 misclassified slides were selected for annotation. Four pathologists (two principal investigators and two residents) participated in the study, each spending approximately two hours on annotation. For each slide, pathologists rated both the attention heatmap and the spectrum-based explanation produced by MorphoXAI along three dimensions—ease of understanding, specificity of diagnostic information, and helpfulness for diagnostic decision-making. In addition, for the spectrum-based explanations, pathologists further evaluated the morphologic consistency between high-contribution patches and their corresponding spectrum clusters.

### 2.8 Validation of MorphoXAI Explanations

#### 2.8.1 Human Evaluation Setup

In the ovarian study, we conducted a blinded human evaluation using MorphoExplainer, involving four pathologists who had not participated in the earlier phenotyping of the spectrum clusters and were blinded to the independent test cohort. During the evaluation, pathologists reviewed slides from the independent cohort together with the model’s predicted subtype probabilities and the corresponding slide-level explanations generated by MorphoXAI. All assessments were performed within the MorphoExplainer interface, and feedback was collected through its built-in scoring panel.

To ensure a balanced and representative assessment, we sampled 48 correctly predicted slides from the independent test set, selecting 12 slides per subtype. All 26 misclassified slides from the independent cohort were also included (Fig. 5b).

The evaluation consists of two parts. The first part compared the impact of attention heatmaps and spectrum-based explanations on pathologists’ ability to understand and verify the model’s predictions. For each slide, pathologists sequentially examined the attention heatmap and the MorphoXAI slide-level explanation and rated each explanation along three dimensions—ease of understanding, specificity of diagnostic information, and helpfulness for diagnostic decision-making (question details and scoring scales are provided in Methods and Supplementary Note 3). In total, 296 rating records were collected for each explanation type. In the second evaluation part, pathologists examined the MorphoXAI slide-level explanations by checking whether the morphology of each high-contribution patch, as observed on the WSI, aligned with the mapped pattern defined in the morphologic spectrum. This process yielded 1,808 assessments (452 spectrum clusters × 4 participants; see Supplementary Table 1 for statistics).

#### 2.8.2 Alignment Between the Morphologic Spectrum and the Model’s Slide-Level Prediction Behavior

We first evaluated the model’s reliance on the morphologic spectrum when predicting unseen slides by analyzing, within the slide-level explanations of the independent cohort, whether the high-contribution patches were enriched in the spectrum patterns corresponding to the predicted subtype. For each slide, we defined the enrichment proportion (*E*) as the fraction of high-contribution patches mapped to the predicted subtype’s patterns in the morphologic spectrum. As Fig. 6a shows, correctly predicted slides exhibited subtype-specific enrichment patterns, with their high-contribution patches predominantly mapped to the histomorphologic clusters associated with their ground-truth subtype. For example, CC, EC, HGSC, and SBT slides showed high mean enrichment (*Ē* = 0.76, 0.66, 0.83, and 0.90, respectively; all *p* < 0.001, Wilcoxon signed-rank test). In contrast, EC slides misclassified as HGSC displayed mixed enrichment patterns. While enrichment within the HGSC clusters was moderate (*Ē* = 0.60), a noticeable fraction of high-contribution patches was also mapped to the EC↔HGSC transitional clusters (*Ē* = 0.36), indicating that the model’s errors were influenced by transitional patterns shared between the two subtypes (see Supplementary Fig. 6 for enrichment patterns at the individual-cluster level).

**Figure 6.**
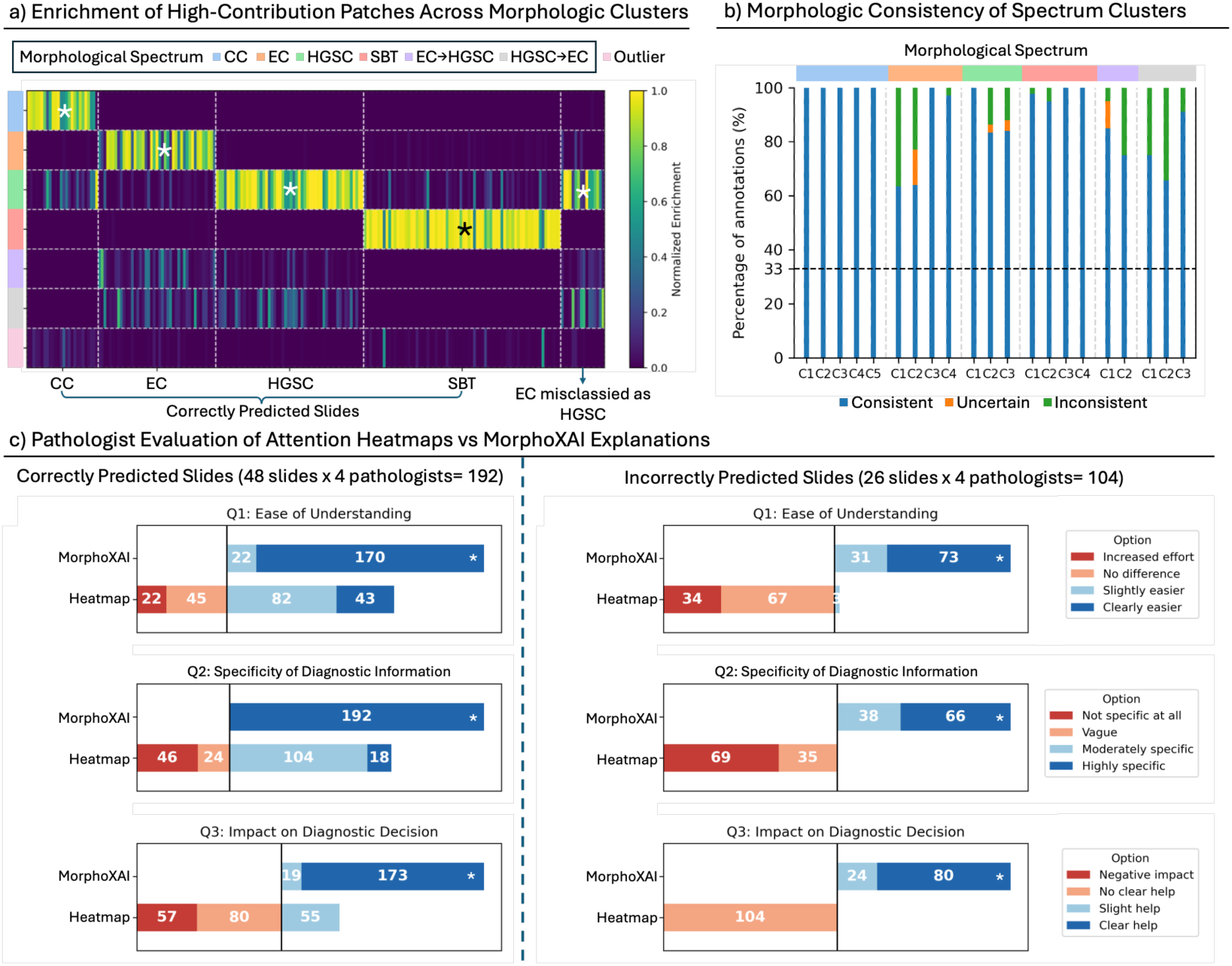
Validation results of MorphoXAI explanations. **a)** Enrichment of high-contribution patches across morphologic clusters. Each column represents a slide from the independent test set, and each row corresponds to a morphologic cluster within the pre-defined spectrum. Each cell (saliency value) indicates the proportion of a slide’s high-contribution patches that were mapped to a given cluster, normalized by the total number of high-contribution patches in that slide. **b)** Morphologic consistency of spectrum clusters. Each bar summarizes pathologists’ annotations assessing the morphologic agreement between mapped high-contribution patches and their assigned spectrum clusters. The three stacked segments within each bar represent the proportions of ratings categorized as consistent, uncertain, and inconsistent, respectively. **c)** Pathologist evaluation comparing attention heatmaps with morphologic spectrum–based interpretations. Asterisks (*) indicate statistically significant results.

Pathologists further examined whether the morphology of each high-contribution patch, as observed on the original WSI, was consistent with the histomorphologic pattern to which it was mapped (Fig. 6b). Many of these patches were judged morphologically consistent with their assigned patterns. The defining patterns of EC and HGSC, as well as the EC↔HGSC transitional patterns showed slightly reduced consistency (≈60–85%) but remained well above the 33% chance baseline, indicating that the MorphoXAI explanations provide morphologically accurate interpretations for most high-contribution patches.

The enrichment analysis shows that, when correctly predicting unseen cases, the model consistently relies on the subtype-defining patterns captured in the morphologic spectrum, whereas in EC cases misclassified as HGSC, it draws on the transitional patterns between these subtypes. This indicates that the model’s prediction behavior on individual slides aligns with how the morphologic spectrum explains both correct and confused classifications at the global level. The pathologists further confirmed this alignment by tracing mapped high-contribution patches back to their morphology on the original WSIs. Together, these findings provide strong support for the morphologic spectrum as a faithful explanation of the model’s predictions, demonstrating the effectiveness of MorphoXAI in interpreting model behavior at both the global and local levels.

#### 2.8.3 Interpretability and Diagnostic Utility Across Diagnostic Scenarios

We next evaluated whether the MorphoXAI slide-level explanations could assist pathologists in understanding and verifying the model’s predictions in diagnostic scenarios, using the attention heatmap as a baseline for comparison (Fig. 6c).

Pathologists reported that the model equipped with MorphoXAI explanations better supported diagnostic decision-making, owing to the explanations being easier to interpret and more informative in revealing specific diagnostic features (all *p* < 0.001; Wilcoxon signed-rank test). Notably, this advantage was even greater for incorrectly predicted slides. To examine how MorphoXAI assists pathologists in understanding and verifying model predictions, we further evaluated its utility across three representative diagnostic scenarios: (i) cases with diagnostic morphology where the model’s prediction was correct, (ii) cases with diagnostic morphology where the model’s prediction was incorrect, and (iii) challenging cases in which morphology alone was insufficient for establishing a diagnosis. For each scenario, we selected a representative case and presented the model prediction, the attention heatmap, and the MorphoXAI explanation to pathologists through MorphoExplainer, and captured their feedback on how each explanation influenced their interpretation.

First, we examined an easy case that exhibited abundant EC defining patterns throughout the slide (Extended Data Fig. 7). The model predicted EC correctly with a high confidence of 98%. Upon reviewing the explanations, pathologists immediately verified the prediction. Both the attention heatmap and the MorphoXAI explanation directed their attention to regions displaying typical EC morphologies, allowing for effortless confirmation of the prediction. However, whereas the heatmap merely highlighted diagnostically relevant areas, MorphoXAI further distinguished the specific histomorphological patterns within these regions—such as squamous metaplasia and glandular patterns—thus providing more granular and clinically specific diagnostic information. This case illustrates that, for easy and correctly predicted slides, both types of explanations are easy for pathologists to understand; however, the MorphoXAI explanations offer more specific diagnostic cues, leading pathologists to perceive them as more helpful for diagnosis, as reflected in the Fig. 6c.

Next, we examined a case that exhibited characteristic histomorphologies of SBT but also contained areas with glandular patterns (Extended Data Fig. 8). The model incorrectly predicted this slide as EC, assigning a probability of 62% to EC and 38% to SBT. When reviewing the EC explanation, pathologists noted that the MorphoXAI explanation highlighted regions of squamous metaplasia and glandular pattern. However, the regions mapped to squamous metaplasia actually corresponded to normal epithelium, immediately raising doubts about the correctness of the prediction. Subsequent inspection of the MorphoXAI explanation for SBT directed the pathologists to two key diagnostic patterns—papillary architecture and simple epithelial layering—which confirmed that the slide was indeed SBT. This also revealed the model’s tendency to misinterpret solid areas of normal epithelium as squamous metaplasia. In contrast, the attention heatmaps for both the SBT and EC predictions broadly highlighted the tumor regions, appearing nearly identical and offering little insight into why the model had made an error. This case illustrates that, in situations of incorrect prediction, the MorphoXAI explanations can expose model errors and thereby help prevent overtrust by pathologists. Moreover, by revealing the rationale behind the misclassification, the MorphoXAI explanation offers valuable insights for model improvement.

Finally, we examined a diagnostically challenging EC case that exhibited overlapping morphologic features of EC and HGSC (Extended Data Fig. 9). The model assigned comparable probabilities to both subtypes (HGSC: 56%, EC: 44%). When reviewing the HGSC explanation, pathologists observed typical HGSC patterns—pleomorphism, high nuclear-to-cytoplasmic ratio, and nuclear hyperchromasia—alongside transitional glandular patterns that morphologically resembled EC. Upon switching to the EC explanation, they again noted EC defining patterns together with transitional morphologies. Based on these findings, the pathologists recognized the slide as a mixed-up case requiring further examination. They also noted that the model-identified transitional regions could serve as valuable candidates for biomarker discovery and help automate the detection of transitional patterns that usually require labor-intensive manual search. In contrast, the attention heatmaps for EC and HGSC highlighted largely overlapping tumor regions, offering limited insight into the rich diagnostic diversity present within the slide. This case illustrates that, for diagnostically challenging slides, MorphoXAI explanations enable collaborative reasoning between the model and pathologists and provides richer diagnostic insights, offering even greater assistance than in easy cases.

In summary, MorphoXAI explanations not only improve the interpretability of model predictions but also demonstrate distinct diagnostic utilities across different diagnostic contexts: reinforcing human–AI trust in correct cases, exposing errors in misclassified cases, and facilitating joint reasoning and deeper diagnostic understanding in complex cases. Together, these results highlight the potential of MorphoXAI in enhancing the transparency of deep learning models in clinical settings.

## 3 Discussion

Deep learning models have achieved strong performance in whole-slide image (WSI) classification, yet their lack of transparency regarding which histomorphologic patterns drive their predictions remains a major barrier to clinical deployment. In this work, we introduce MorphoXAI, a human-in-the-loop explanation framework that reveals the histomorphologic patterns a model has learned to distinguish WSI classes at the global level and the specific patterns it relies on when predicting an individual slide at the local level. By unifying global and local explanations and grounding the explanations in expert-interpreted morphology collected through a human-in-the-loop process, MorphoXAI enhances the transparency of deep learning models and supports more effective pathologist–AI collaboration in diagnostic settings. Our evaluation on ovarian tumor subtyping demonstrates the effectiveness of this framework, showing that MorphoXAI provides meaningful insight into model behavior and assists pathologists in understanding and verifying model predictions.

Equipping deep learning models with MorphoXAI explanations could strengthen human–AI collaboration by enabling models to provide clinically meaningful diagnostic support and allowing pathologists’ feedback to inform model refinement (Extended Data Fig. 7–9). For pathologists, unlike attention heatmaps that merely highlight spatial saliency, MorphoXAI explanations localize and clarify the morphological evidence underlying model predictions, making the model’s predictive rationale not only visually intuitive but also semantically clear, thereby improving pathologists’ ability to interpret and verify model outputs and fostering greater trust in the AI system. MorphoXAI could also enable pathologists to contribute to refining deep learning systems. Through MorphoExplainer, experts can actively identify discrepancies between the model’s reasoning and established diagnostic knowledge (Extended Data Fig. 8), pinpointing patterns that are diagnostically irrelevant, misleading, or overlooked. Such feedback provides meaningful guidance for model optimization, improving not only predictive performance but also the interpretability and reliability of the system. Before clinical deployment, this framework can also serve as a practical tool for auditing a model’s learned morphological knowledge. Such assessment complements quantitative validation by adding an additional layer of quality control, ensuring that model predictions are not only accurate but also grounded in credible morphologic evidence.

Under research settings, MorphoXAI could be used to identify transitional morphologic patterns and to characterize the morphologic spectra underlying disease evolution (Extended Data Fig. 9). Characterizing the morphologic changes that occur across disease stages is central to understanding pathogenic mechanisms and establishing diagnostic criteria^42^. Traditionally, such spectra are delineated manually by pathologists through the curation of representative cases and the qualitative description of their morphologic features—a process inherently limited by human effort and data scale^43–46^. MorphoXAI offers a data-driven alternative for this research goal. By leveraging growing archives of WSIs with subtype annotations, one can train deep learning models and, through the MorphoXAI framework, obtain a model-derived morphologic spectrum. This approach could improve the efficiency of constructing disease-relevant morphologic continua, support biomarker discovery, and inform mechanistic studies of disease.

There are several limitations to our study. First, our study focuses on tumor subtyping as the representative WSI-level classification task and uses an ABMIL architecture as the model backbone. However, deep learning for WSI classification encompasses a broader range of predictive objectives—such as patient survival, treatment response, and genetic alterations—and includes diverse model architectures, including graph-based^47^ and transformer-based^48^ frameworks. Future work could extend the MorphoXAI framework to these additional prediction tasks and model architectures to further assess its generalizability. Second, our pathologist-centric evaluation involved a small but highly qualified group of board-certified pathologists, prioritizing in-depth analysis and expert-level feedback. Although the results were encouraging and the sample size was comparable to prior XAI studies in pathology^49–53^, future evaluations involving larger and more diverse groups of users could more comprehensively validate the effectiveness of MorphoXAI explanations in improving model interpretability and their utility in diagnostic decision-making.

On the methodological side, several aspects of our framework warrant improvement. First, the selection of thresholds for sample stratification and the determination of optimal cluster numbers currently rely on empirical choices; developing more objective or automated strategies could enhance the robustness and reproducibility of the analysis. Second, our approach benefits from expert interpretation of morphology. Although the interactive tools we developed, MorphoCA and MorphoExplainer, facilitate the efficient collection of expert feedback and allow pathologists to trace model predictions back to the original WSIs, they are currently implemented within the QuPath platform. Further development is needed to improve cross-platform compatibility and integration, thereby accommodating a wider range of user environments and workflows.

While many existing XAI techniques are fully data-driven or incorporate pathology priors only through predefined expert concepts, MorphoXAI brings domain expertise into the explanation process through human-in-the-loop interactions. By enabling human experts to access, investigate, and interpret model prediction behavior on the original WSI, MorphoXAI grounds explanations in expert-validated morphology, offering a complementary perspective for achieving transparency and interpretability in medical AI systems. The application of MorphoXAI in future XAI studies could yield meaningful insights into how AI models operate and help advance the transparent and trustworthy deployment of medical AI systems.

## 4 Methods

### 4.1 Ovarian Datasets

The study included two ovarian tumor cohorts collected at Mayo Clinic and prepared using standardized histopathologic workflows. All specimens were obtained from surgical resections and were anonymized prior to analysis. The study was approved by the Mayo Clinic Institutional Review Board.

The training cohort consisted of ovarian tumor resections collected between 2022 and 2024. All specimens were processed with routine hematoxylin and eosin (H&E) staining. Slides were digitized using a Leica Aperio GT450 scanner at 40× magnification (0.25 µm/pixel) and stored in .svs format. Quality control was performed through double review by board-certified pathologists, and slides were excluded if they contained mixed histologic components, insufficient tissue, or prior treatment effects. Subtype confirmation was supported by immunohistochemical markers, including Napsin A and HNF-1β for Clear Cell Carcinoma (CC); WT1, p53, and p16 for High-grade Serous Carcinoma; and ER, PR, and p16 for Endometrioid Carcinoma. The final training set comprised 68 patients (602 WSIs) representing four histologic subtypes: Serous Borderline Tumor (21 patients, 134 WSIs), High-grade Serous Carcinoma (18 patients, 230 WSIs), Clear Cell Carcinoma (11 patients, 101 WSIs), and Endometrioid Carcinoma (18 patients, 137 WSIs).

The independent test cohort included ovarian tumor resections collected between 2024 and 2025, with no patient overlap with the training cohort. All specimens followed identical processing, staining, scanning, and quality control protocols as the training cohort, with the same immunohistochemical markers used for subtype confirmation. This cohort included 31 patients (221 WSIs): Serous Borderline Tumor (11 patients, 77 WSIs), High-grade Serous Carcinoma (8 patients, 56 WSIs), Clear Cell Carcinoma (4 patients, 27 WSIs), and Endometrioid Carcinoma (8 patients, 61 WSIs).

### 4.2 CLAM Model Architecture

We adopted CLAM for whole-slide image classification and applied our explanation framework upon it, which has been widely applied in WSI classification across multiple disease types.

CLAM operates as follows: first, each WSI is partitioned into *N* non-overlapping image patches, each represented by a high-dimensional embedding vector *h*_*k*_ ∈ *R*^*d*^, obtained from an encoder. For each class *c*, which in this study corresponds to one histologic subtype, CLAM computes class-specific attention weights *a*_*k*,*c*_ for each patch *k* via an attention gate mechanism:

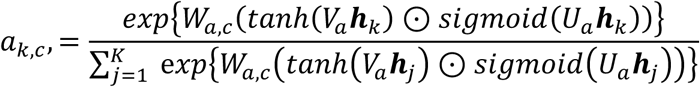

where *W_a,c_*, *V_a_* and *U_a_* are learnable parameters, t*anh*(·) and *sigmoid*(·) denote element-wise hyperbolic tangent and sigmoid activation functions, respectively, and ⊙ indicates element-wise multiplication. *a*_k,c_ quantifies the importance of each patch for predicting *c*. Next, the attention weights are used to aggregate patch embeddings into a class-specific slide-level representation ***z***_c_:

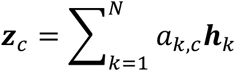

Subsequently, slide-level logits *S_slide,c_* for class *c* are computed via a linear classifier parameterized by weight vector *W_c_*:

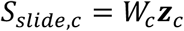

Finally, the logits vector 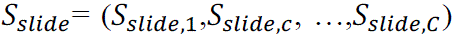 is converted into probabilities using the softmax function, yielding the predicted class probability distribution at the slide level.

In this study, we replaced the original CNN-based encoder used by CLAM with CONCH^33^, a pretrained encoder specifically optimized for histopathological images, to enhance the representation of patch-level morphologic features.

### 4.3 Model Training and Testing

This section outlines how the full-data model was trained and how the multiple training–testing runs required for prediction-stability–based sample stratification were carried out.

The full-data model was trained on the entire training cohort dataset. Hyperparameters were first selected through 10-fold cross-validation on the training set. Four candidate configurations were evaluated, and the configuration achieving the highest mean macro-average AUC across folds was subsequently adopted. The final model was optimized using the Adam optimizer with an initial learning rate of 1 × 10^01^ and a weight decay of 1 × 10^02^. The learning rate was decayed by a factor of 0.1 at epochs 5, 15, and 30 following a step decay schedule. A cross-entropy loss was used for optimization, and training was conducted for 150 epochs. Dropout probabilities were set to 0.25 for the attention layer and 0.5 for the fully connected layer. Each iteration used a batch size of one slide, containing all patch-level feature vectors from a single WSI, randomly sampled from the training dataset. The complete hyperparameter configuration is listed in Supplementary Table 2.

To construct the dataset splits for the training and testing runs required for prediction-stability–based sample stratification (Supplementary Fig. 2a), the ovarian cancer cohort was randomly shuffled at the patient level (based on Patient ID) five times using different random seeds, generating five distinct patient permutations. In each permutation, patients—together with all their slides—were sequentially assigned to ten partitions in a round-robin manner (e.g., the first patient to partition 0, the second to partition 1, …, the tenth to partition 9, then the eleventh back to partition 0, and so on) until all patients were assigned. All slides from the same patient were kept within a single partition to avoid data leakage. The ten partitions were then allocated to training, validation, and testing sets in a 6:2:2 ratio. Consequently, each partition was included in the test set twice within a permutation, such that every patient (and thus every slide) was evaluated ten times across the five permutations.

For each data split, a CLAM model was trained, validated, and tested independently using the same set of hyperparameters (Supplementary Fig. 2b). Unlike the full-data model, these models adopted an early stopping strategy with a patience of 20 epochs, based on validation loss, to prevent overfitting and reduce redundant computation during repeated training. This process yielded ten trained models (one per split) for each patient permutation, resulting in a total of 50 trained models across the five permutations. Performance metrics (e.g., AUC) obtained from each test run were averaged across all five permutations to produce the final performance estimate (Supplementary Fig. 2c).

### 4.4 Contribution Score Calculation and Patch Clustering

In this study, rather than using the raw attention weights *a*_!,$_ as the measure of patch importance, we adopt a logit-based contribution score to quantify each patch’s influence on the slide-level prediction logits. Building upon the formulation defined in Section 4.2, the slide-level logit for a class *c* is computed as follows:

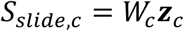

where ***z**_c_* represents the slide-level embedding for class *c*, computed as a weighted sum of patch embeddings ***h**_k_*:

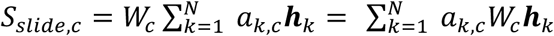

By expanding and rearranging the terms, the slide-level logit can be represented explicitly as a sum of individual patch contributions, where the contribution from each patch *k* to class *c* is defined as follows:

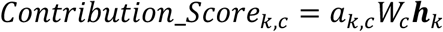

The logit-based contribution score quantifies each patch’s signed influence on the class logit, capturing whether a patch positively or negatively drives the model’s prediction. Furthermore, since the slide-level logit is the sum of these individual contributions, patches can be directly ranked by their impact on the final decision, with the top-ranking patches designated as high-contribution patches.

To identify common histomorphological patterns among the high-contribution patches, we employed an unsupervised clustering workflow based on patch embeddings generated by CONCH. First, all patch embeddings were compressed into 2000 representative micro-clusters using K-means clustering implemented via the FAISS library^54^, ensuring computational efficiency and reducing redundancy. Next, a consensus clustering approach was employed to determine stable phenotypic clusters. Specifically, K-means clustering was iteratively performed on randomly sampled subsets containing 80% of the micro-clusters, with candidate cluster numbers *k* ∈ {2, …, 8}. Each candidate value of *k* was tested through 30 independent subsampling–clustering iterations, generating a consensus matrix in which each element reflected the frequency with which two micro-clusters were assigned to the same cluster across iterations. The optimal cluster number was selected by maximizing the cophenetic correlation coefficient (COPH), providing the highest clustering stability. Finally, hierarchical clustering with complete linkage was applied to the original set of patch embeddings with this optimal cluster number, yielding the final set of patch clusters for downstream analyses.

The consensus matrices and COPH used to determine the optimal number of clusters are shown in Supplementary Figures 3–5. For most groups, the optimal *k* was selected by maximizing the COPH value. For the Consistently Correct groups of SBT, EC, and HGSC, although the highest COPH values suggested *k* = 2, we selected slightly larger cluster numbers (*k* = 4 for SBT and EC, and *k* = 3 for HGSC) based on pathological domain knowledge that more than two core morphologic phenotypes are commonly recognized for these subtypes in clinical practice.

### 4.5 High-Contribution Patch Extraction and Mapping for Generating Slide-level Explanation

To generate slide-level explanation, the full-data model was applied to the test cohort, from which high-contribution patches were extracted and subsequently mapped onto the predefined morphologic spectrum. The selection of high-contribution patches followed the same criterion used in constructing the morphologic spectrum: for each subtype prediction, the top-ranked patches whose cumulative contribution scores accounted for 90% of the total slide-level logit were retained.

Spectrum mapping was performed based on patch embeddings encoded by CONCH. First, for each cluster from the morphologic spectrum, we computed the mean vector (µ) and shrinkage covariance matrix (Σ) from the embeddings of all patches assigned to that cluster in the training cohort, representing its center and shape in the embedding space. We next calculated the Mahalanobis distance d(x, µ, Σ) of all the patches to the cluster center, defining the 95th percentile of within-cluster distances (r_32_) as the boundary threshold for that cluster. During mapping, for each high-contribution patch from the independent test set, we computed its Mahalanobis distances to all cluster centers and assigned it to the cluster with the minimal distance. If this distance exceeded the corresponding r_32_, the patch was labeled as an outlier.

For high-contribution patches supporting EC or HGSC predictions, we applied an additional transitional phenotype detection procedure. Specifically, if a patch from the test cohort was initially assigned to an EC or HGSC core cluster, its local neighborhood in the embedding space was identified by performing a k-nearest neighbors (k-NN) search among patches from the training cohort. The fraction of transitional samples among its k-nearest neighbors (p_trans_) was then computed. We further computed the global transitional baseline proportion (p_trans,base_) defined as the overall fraction of transitional patches among all EC, HGSC, EC→HGSC, and HGSC→EC patches in the training cohort. When p_trans_ > p_trans,base_, the patch from the test cohort was identified as transitional; otherwise, it remained assigned to its original core cluster.

The specific mapping transitional cluster was determined through a weighted voting scheme among the nearest transitional neighbors. Each neighbor patch *j* was assigned an inverse-distance weight:

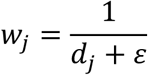

where *d_j_* denotes the embedding distance between the current patch and neighbor *j* and ε is a small constant to avoid division by zero. The weights were then grouped by the transitional cluster label *t* of each neighbor, where *t* represents one of the EC→HGSC or HGSC→EC clusters appearing among the nearest neighbors. The weights within each group were summed to obtain a cumulative score for each candidate cluster:

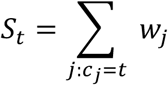

Here, *c_j_* denotes the transitional cluster label of neighbor *j*. Finally, the patch was assigned to the transitional cluster with the highest cumulative score.

### 4.6 Pathologist Annotation of Spectrum Clusters

In the step of cluster phenotyping, board-certified pathologists annotated the morphological characteristics of clusters in the morphologic spectrum. For every cluster, pathologists were asked to answer four structured questions:

- **Represented Phenotype**: What histological or cytological phenotype (i.e. histomorphological pattern) consistently predominates within the majority of patches in this cluster? Pathologists are asked to respond with the name(s) of established phenotype(s) (e.g., “glandular or papillary architecture”). Given individual patches may include artifacts or noise, annotations should focus on the phenotype(s) consistently represented across the entire cluster. If uncertain, pathologists should indicate “N/C” (Not Certain).
- **Subtype-defining Phenotype**: Are the represented phenotype(s) essential diagnostic features for identifying the subtype of the corresponding WSI? Pathologists are asked to answer “Yes”, “No”, or “N/C”.
- **Detailed Morphological Descriptions (Intra-cluster Features)**: List and briefly describe secondary morphological characteristics consistently appearing across most patches in the cluster (e.g., “nuclei of irregular size and shape,” “marked stromal fibrosis”). These detailed annotations provide deeper insights into the cluster’s morphological characteristics. If uncertain, pathologists should indicate “N/C”.
- **Inter-cluster Morphological Differences**: Compared with other clusters representing the same phenotype(s), what subtle morphological features distinguish this cluster? Pathologists are asked to provide 2–3 distinguishing traits (e.g., “Cluster A contains more stromal tissue compared to other adenomatous clusters”). If uncertain, pathologists should indicate “N/C”.

Before annotation, a shared morphologic vocabulary was developed to ensure consistency among participating pathologists. The vocabulary contained a list of keywords describing representative phenotypes and morphological characteristics associated with the four subtypes (SBT, EC, HGSC, and CC). During annotation, pathologists were encouraged to use terms from this vocabulary whenever applicable.

To clarify the histomorphological patterns represented by each cluster, we performed keyword enrichment analysis on the pathologists’ annotations. For each cluster, morphological keywords were extracted from the pathologists’ responses to the “Represented Phenotype” and “Detailed Morphological Descriptions” questions using the shared morphologic vocabulary. The frequency of each keyword was then quantified across clusters to identify enriched morphologic features. Subsequently, hierarchical clustering with Ward’s method and Euclidean distance was performed independently on both keywords (rows) and clusters (columns), and the reordered data matrix was visualized as a heatmap (Fig. 4a and Extended Data Fig. 2-6a).

### 4.7 Pathologist-centric Interpretability Evaluation

To evaluate the interpretability and diagnostic usefulness of MorphoXAI explanations, board-certified pathologists were invited to compare the spectrum-based explanations and the attention heatmaps (Fig. 5b, 5c). Three structured questions were designed to evaluate pathologists’ comprehension and utilization of model explanations:

- **Ease of understanding**: compared with finding diagnostic features on your own, how did this explanation affect the time and effort you needed to understand the slide? Scores: 0 = Increased effort, 1 = No difference, 2 = Slightly easier, 3 = Clearly easier.
- **Specificity of diagnostic information**: how specific or detailed is the diagnostic information provided by this explanation? Scores: 0 = Not specific at all, 1 = Vague, 2 = Moderately specific, 3 = Highly specific.
- **Impact on diagnostic decision**: to what extent does this explanation help you make or adjust your diagnostic decision? Scores: 0 = Negative impact, 1 = No clear help, 2 = Slight help, 3 = Clear help.

The full questionnaire is provided in Supplementary Note 3. The collected scores were used to quantitatively compare interpretability between the spectrum-based and heatmap explanations (Fig. 6c).

### 4.8 QuPath Plugin Development

Two QuPath plugins were developed to help pathologists review model outputs and collect annotations:

- **MorphoCA** – designed to assist pathologists in reviewing the morphologic spectrum. The plugin overlays clustered high-contribution patches on WSIs, allowing pathologists to explore individual clusters, freely zoom into regions of interest, and record structured annotations for each cluster.
- **MorphoExplainer** – developed for interactive investigation of the model’s slide-level predictions, the spectrum-based explanations, and the attention heatmap within the context of the original WSI. It allows pathologists to switch between explanation modes and subtype predictions, inspect regions at multiple zoom levels, and record feedback through dedicated scoring panels.

Both plugins were implemented in Java using the QuPath SDK^55^ (version 0.6.0) and the QuPath extension framework^56^, and are released together with this paper. The usage instructions are provided in Supplementary Notes 1 and 2 respectively.

### 4.9 Ethics Statement

The study was approved by Mayo Clinic. The ethical approval number is 18-002752.

### 4.10 Data Availability

The datasets used in this study were collected at Mayo Clinic and are not publicly available due to patient privacy considerations and institutional restrictions. Access to these data may be granted by the corresponding author upon reasonable request, subject to approval by the Mayo Clinic Institutional Review Board and completion of appropriate data use agreements.

### 4.11 Code Availability

All code for the MorphoXAI and the QuPath extensions are available at GitHub: https://github.com/dimi-lab/MorphoSpectrum-MIL. Usage instructions and dependencies are provided in the repository README.

### 4.12 Author contributions

P.L.L. and R.F.G. conceived the study. P.L.L. and C.W. designed the experiments. P.L.L. performed the experimental analysis. Y.J.H., R.K., and W.Y. collected and curated the in-house datasets. Y.J.H., R.K., Y.W., and R.F.G. performed the annotation. P.L.L. developed MorphoXAI. B.C.N. tested the code of MorphoXAI. P.L.L. and C.W. prepared the manuscript. Y.Z., N.C., S.J.W., E.L.G., and W.H. assisted with manuscript editing. C.W. supervised the research.

## Figures

**Extended Data Figure 1.**
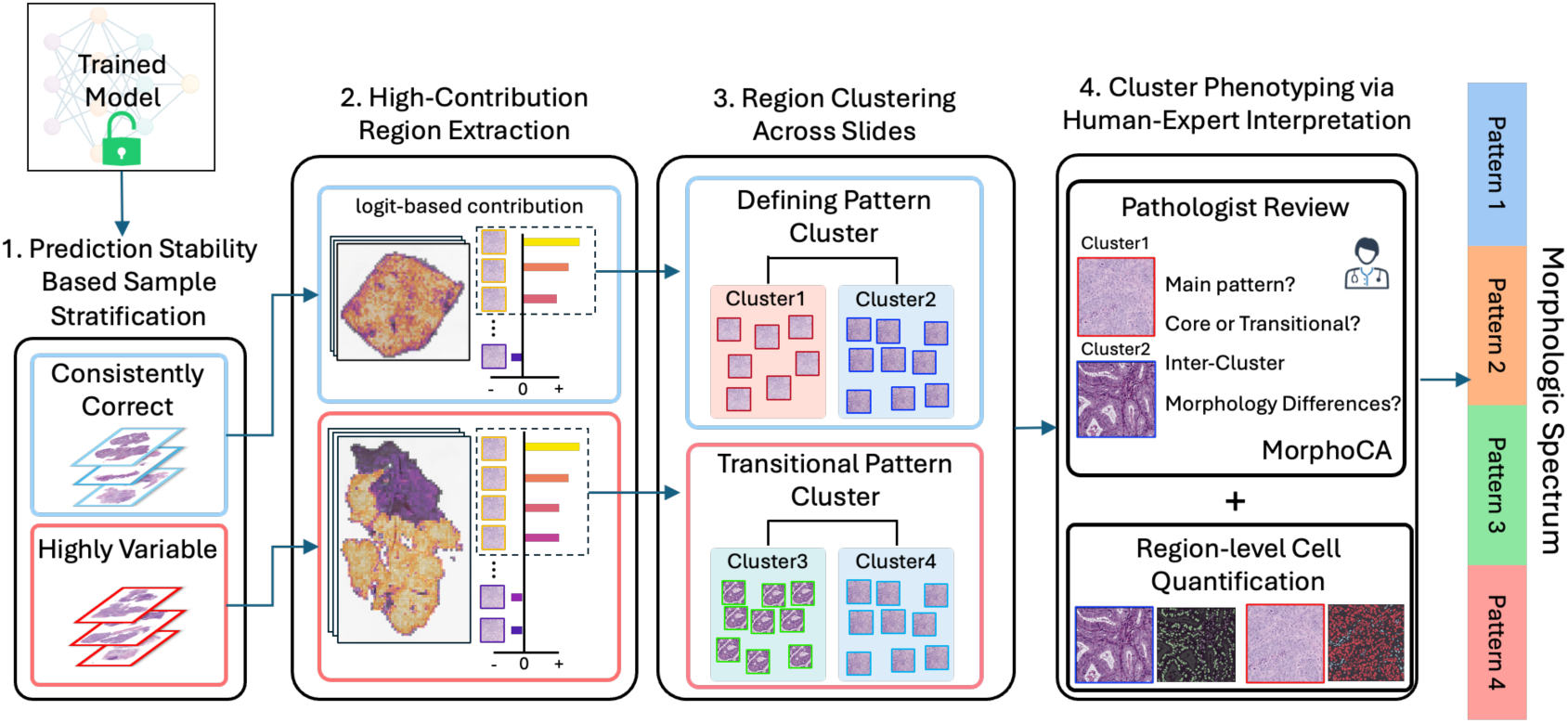
Detailed illustration of morphologic spectrum construction.

**Extended Data Figure 2.**
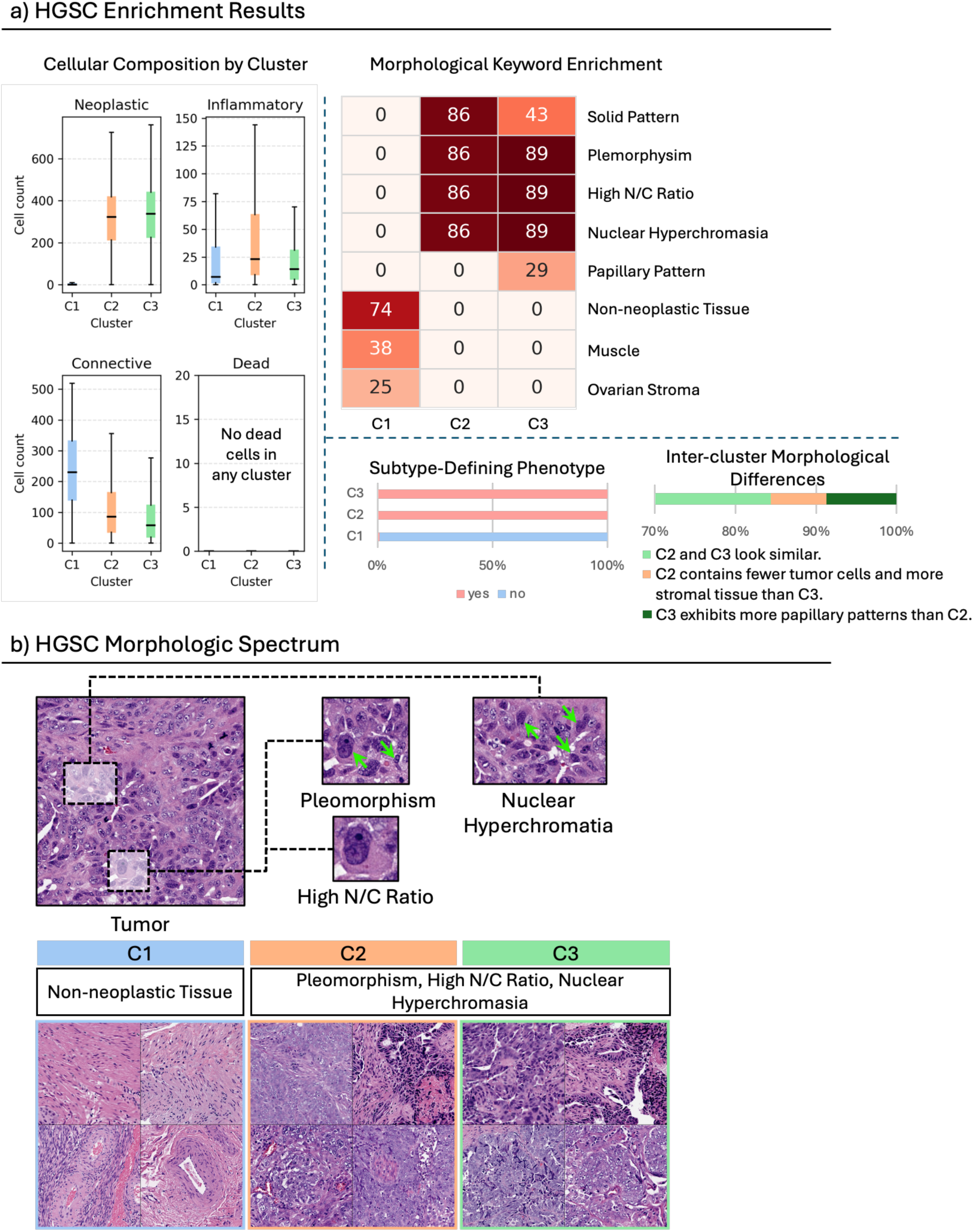
Class-defining patterns for Ovarian High-grade Serous Carcinoma (HGSC). **a)** Enrichment analyses for clusters C1–C3. **b)** Representative image galleries illustrating the histomorphological patterns associated with each cluster. Green arrows highlight specific morphology corresponding to the textual descriptions.

**Extended Data Figure 3.**
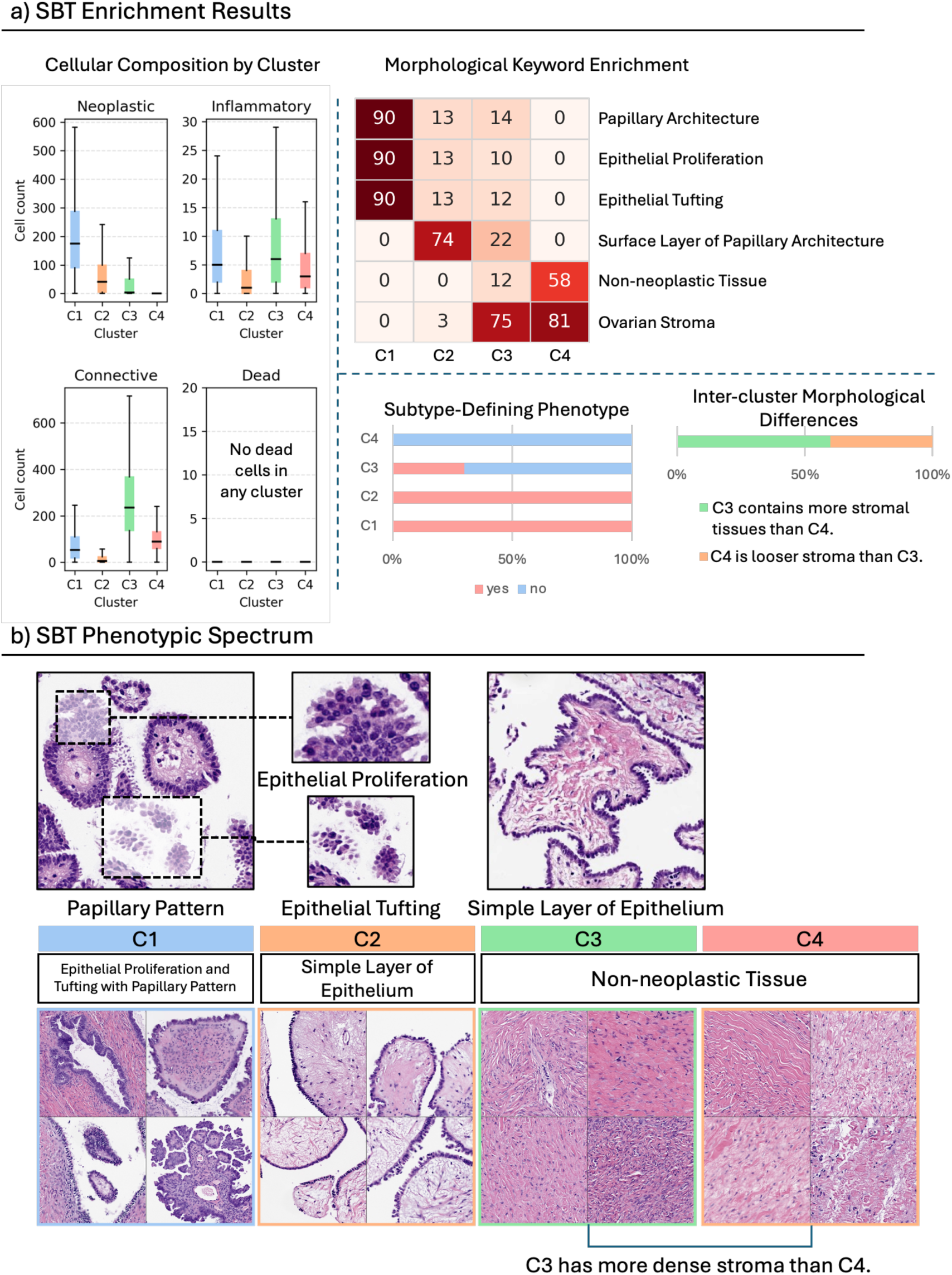
Class-defining patterns for Ovarian Serous Borderline Tumor (SBT). **a)** Enrichment analyses for clusters C1–C4. **b)** Representative image galleries illustrating the histomorphological patterns associated with each cluster.

**Extended Data Figure 4.**
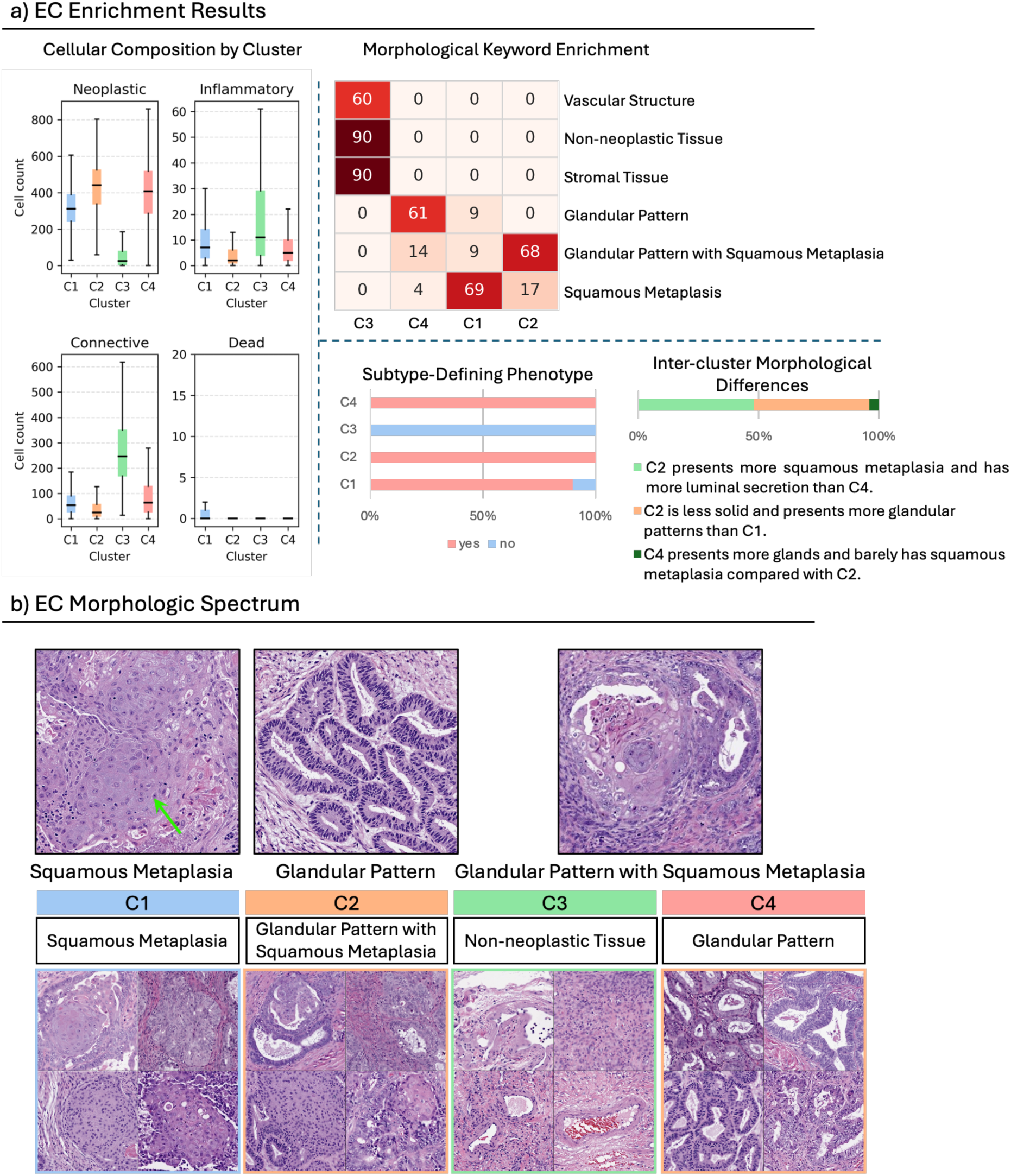
Class-defining patterns for Ovarian Endometrioid Carcinoma (EC). **a)** Enrichment analyses for clusters C1–C4. **b)** Representative image galleries illustrating the histomorphological patterns associated with each cluster. Green arrows highlight specific morphology corresponding to the textual descriptions.

**Extended Data Figure 5.**
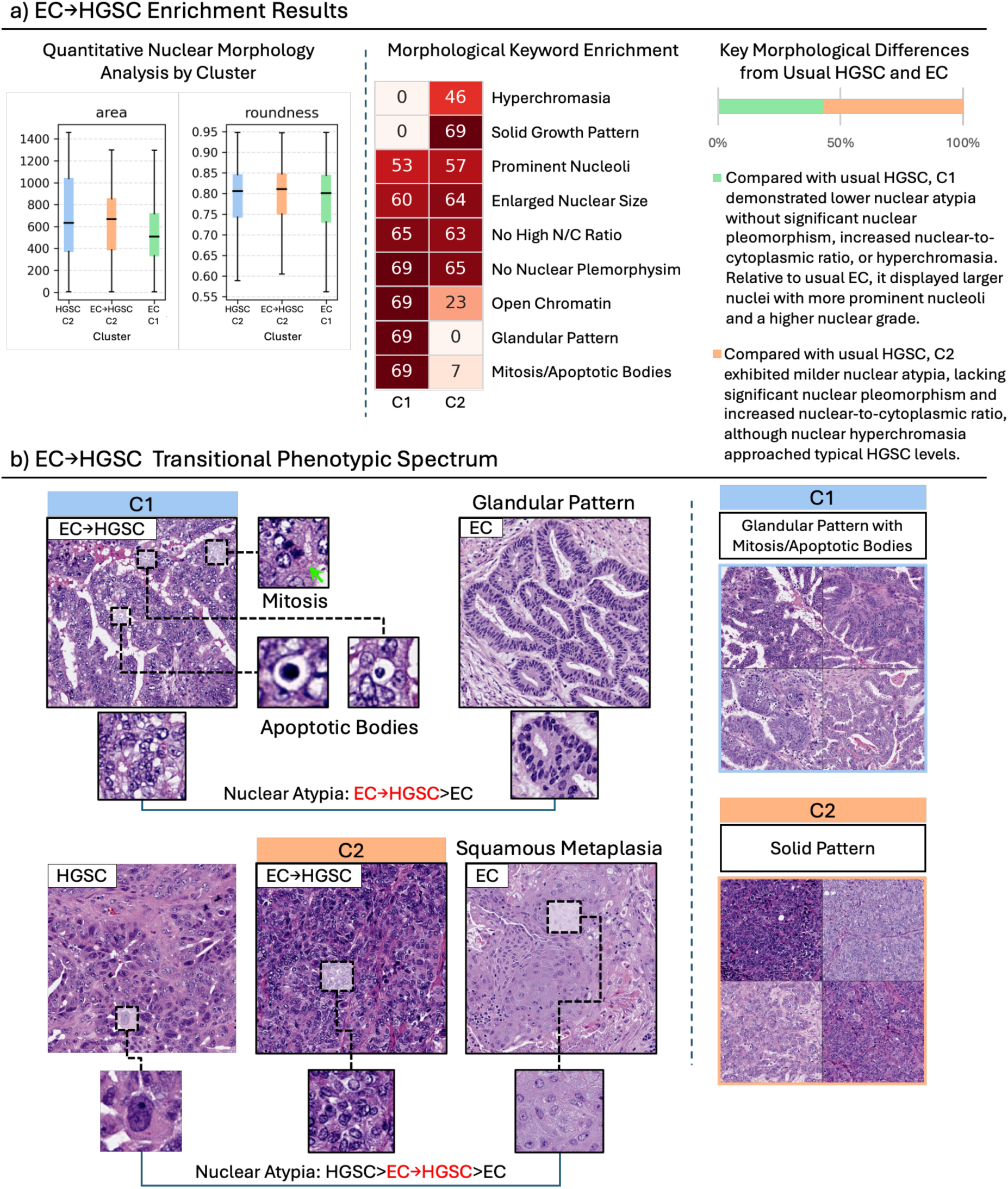
Transitional patterns derived from highly variable Endometrioid Carcinoma slides frequently misclassified as High-grade Serous Carcinoma (EC→HGSC). **a)** Enrichment analyses for clusters C1 and C2. **b)** Representative image galleries illustrating the histomorphological patterns associated with each cluster. Green arrows highlight specific morphology corresponding to the textual descriptions.

**Extended Data Figure 6.**
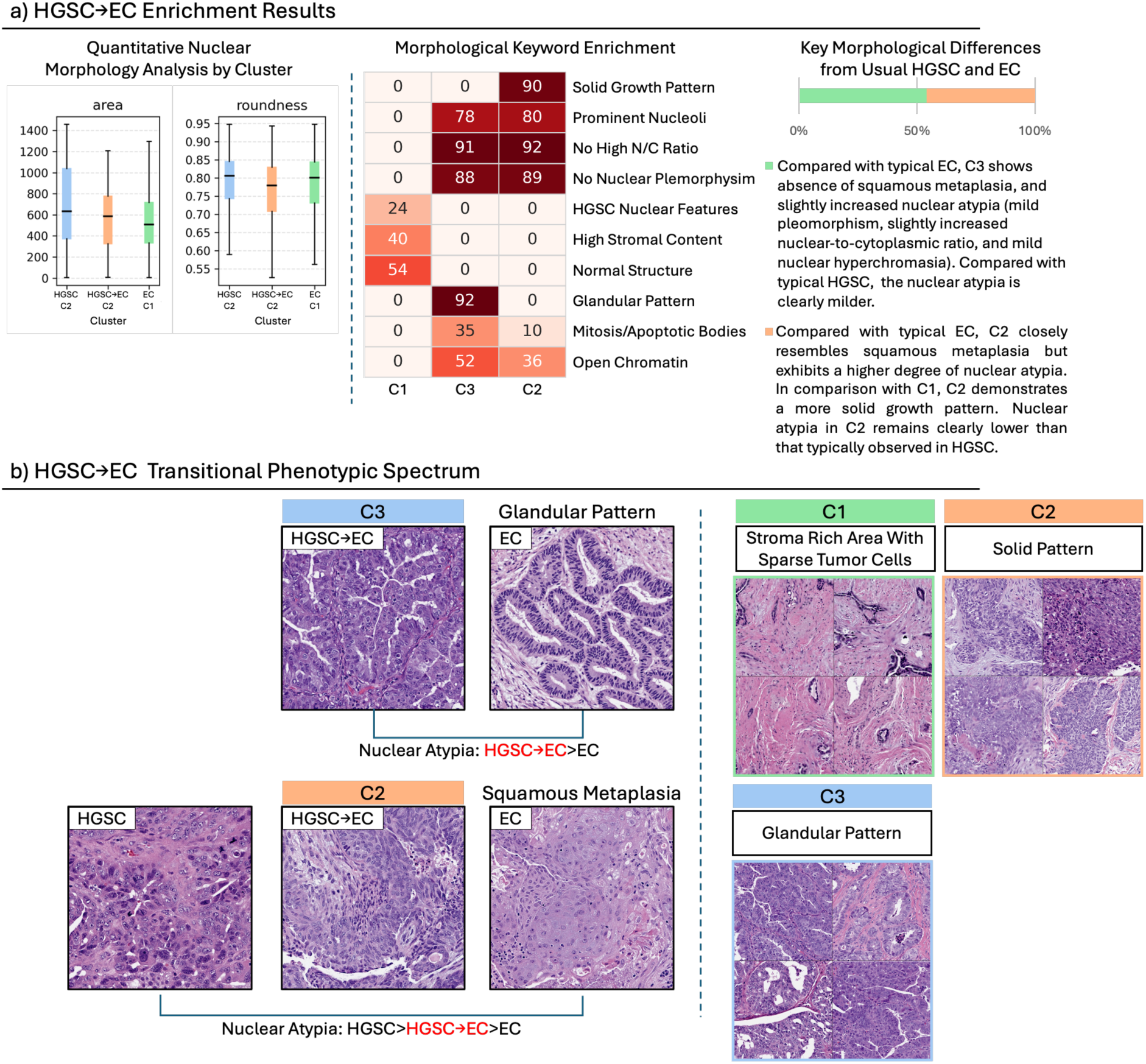
Transitional patterns derived from highly variable High-grade Serous Carcinoma slides frequently misclassified as Endometrioid Carcinoma (HGSC→EC). **a)** Enrichment analyses for clusters C1–C3. **b)** Representative image galleries illustrating the histomorphological patterns associated with each cluster.

**Extended Data Figure 7.**
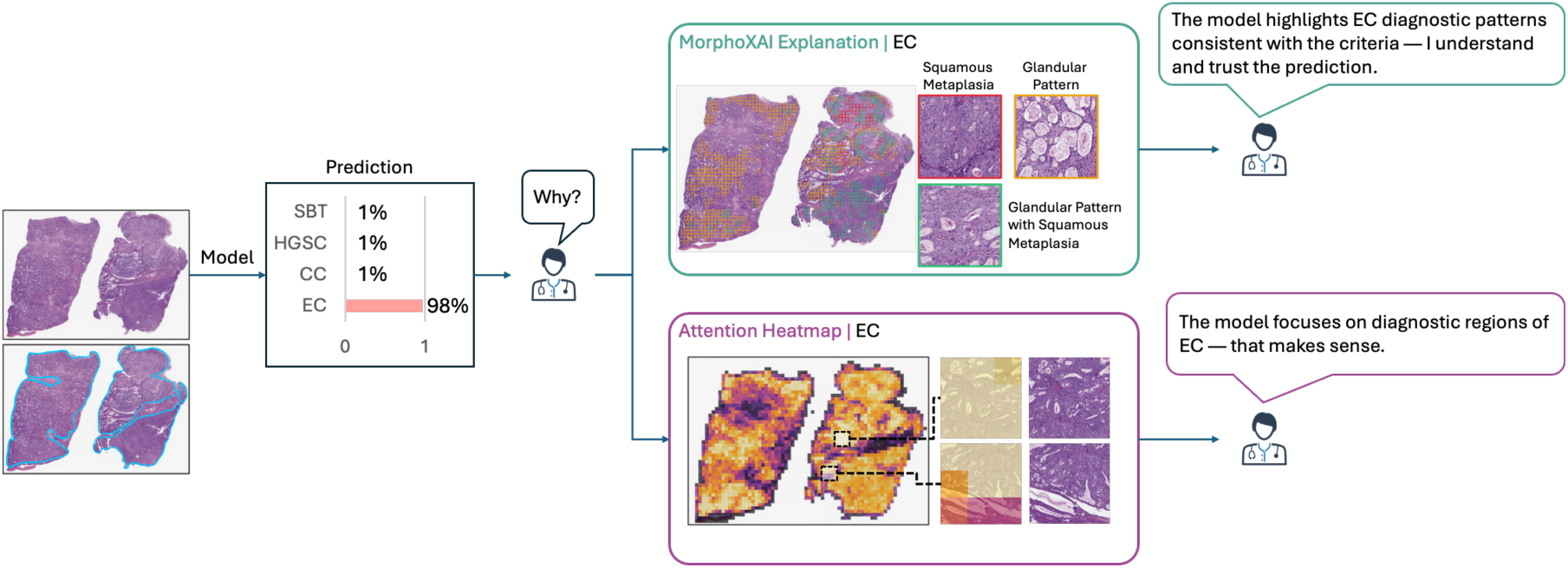
Easy case: comparison of the MorphoXAI explanation and the attention heatmap for a correct and confident EC prediction. The slide exhibited abundant EC morphologies, and the model predicted EC correctly with high confidence (98%). Both the MorphoXAI explanation and the attention heatmap directed pathologists to regions with canonical EC features, but the MorphoXAI explanation further distinguished squamous metaplasia and glandular patterns, providing more specific diagnostic information. For reference, the original slide is accompanied by an annotated image in which the tumor tissue regions are highlighted with blue contours.

**Extended Data Figure 8.**
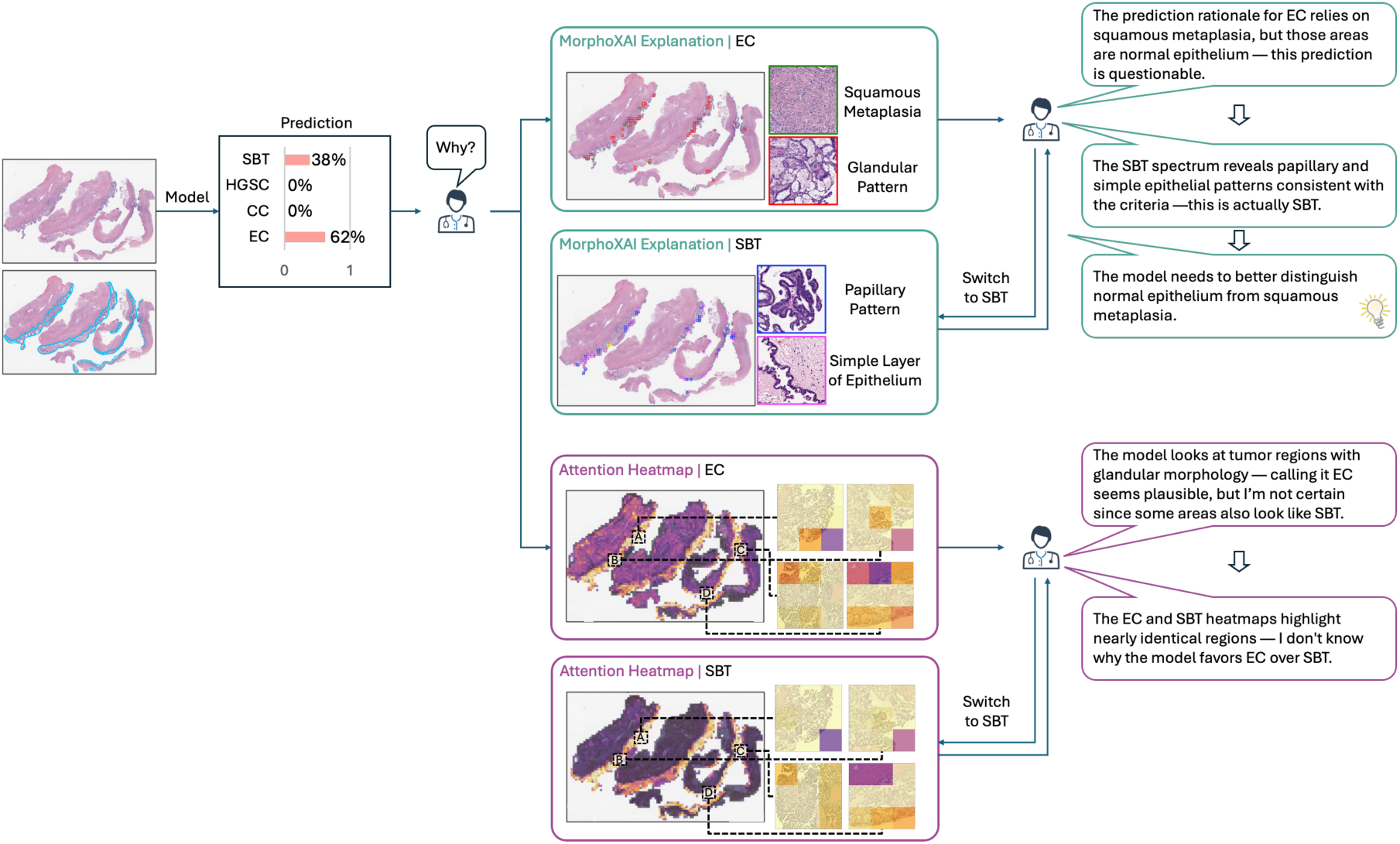
Misclassified case: comparison of the MorphoXAI explanation and the attention heatmap for an SBT slide predicted as EC. The model incorrectly predicted this SBT slide as EC (62% EC, 38% SBT). The MorphoXAI explanation of EC highlighted squamous metaplasia and glandular patterns, but pathologists recognized the former as normal epithelium, revealing the cause of misclassification. In contrast, the MorphoXAI explanation of SBT correctly emphasized papillary and simple epithelial patterns, while attention heatmaps for both subtypes appeared indistinguishable. For reference, the original slide is accompanied by an annotated image in which the tumor tissue regions are highlighted with blue contours.

**Extended Data Figure 9.**
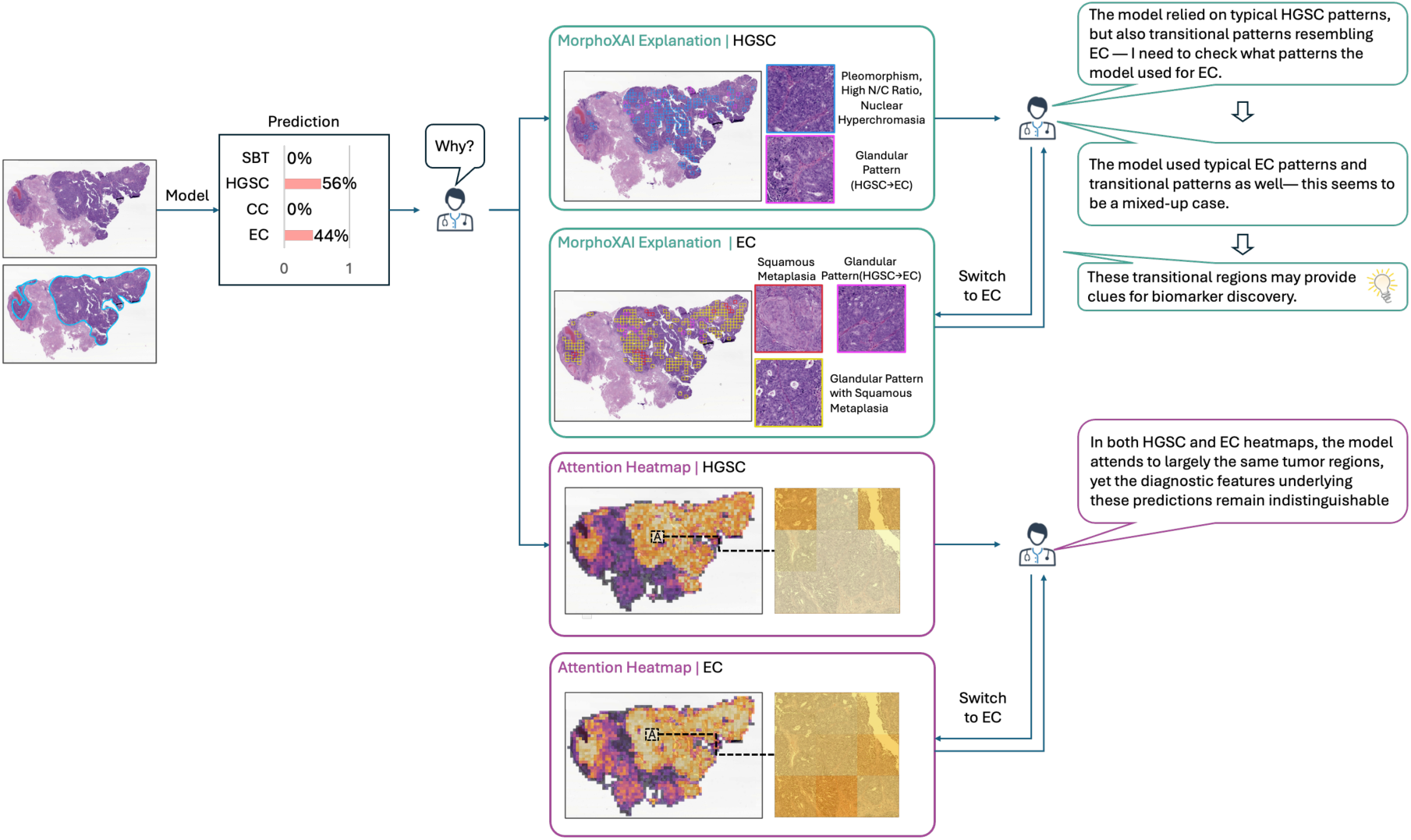
Mixed-up case: comparison of the MorphoXAI explanation and the attention heatmap for a diagnostically complex slide showing overlapping EC and HGSC features. The slide contained both EC– and HGSC-like regions, and the model assigned comparable probabilities to EC (44%) and HGSC (56%). The MorphoXAI explanation revealed transitional glandular patterns, prompting pathologists to classify it as a mixed-up case. These transitional features could serve as valuable candidates for biomarker discovery. For reference, the original slide is accompanied by an annotated image in which the tumor tissue regions are highlighted with blue contours.

