## Supplementary Materials for "A human-in-the-loop explanation framework for morphologically transparent AI predictions from whole-slide images"

Supplementary Table 1: Distribution of morphologic-spectrum clusters and pathologist consistency evaluations

| Cluster | Count* | Consistent | Inconsistent | Uncertain |
|---|---|---|---|---|
| Border_C1 | 34 | 133 | 3 | 0 |
| Border_C2 | 44 | 167 | 9 | 0 |
| Border_C3 | 2 | 8 | 0 | 0 |
| Border_C4 | 1 | 4 | 0 | 0 |
| Clear_C1 | 1 | 4 | 0 | 0 |
| Clear_C2 | 1 | 4 | 0 | 0 |
| Clear_C3 | 1 | 4 | 0 | 0 |
| Clear_C4 | 19 | 76 | 0 | 0 |
| Clear_C5 | 29 | 116 | 0 | 0 |
| Endo_C1 | 28 | 71 | 41 | 0 |
| Endo_C2 | 59 | 151 | 54 | 31 |
| Endo_C3 | 2 | 8 | 0 | 0 |
| Endo_C4 | 59 | 229 | 7 | 0 |
| High_C1 | 3 | 12 | 0 | 0 |
| High_C2 | 33 | 110 | 18 | 4 |
| High_C3 | 89 | 299 | 43 | 14 |
| High_to_Endo_C1 | 2 | 6 | 2 | 0 |
| High_to_Endo_C2 | 8 | 21 | 11 | 0 |
| High_to_Endo_C3 | 31 | 113 | 11 | 0 |
| Endo_to_High_C1 | 5 | 17 | 1 | 2 |
| Endo_to_High_C2 | 1 | 3 | 1 | 0 |

*The total number of times high-contribution patches in the MorphoXAI slide-level explanations of the sampled independent test set used for human evaluation were mapped to that spectrum cluster.

Supplementary Table 2: Model Training Hyperparameters

| Hyperparameter | Value |
|---|---|
| Maximum Epochs | 150 |
| Learning Rate (LR) | 1e-4 |
| Early Stopping Patience* | 20 epochs |
| LR Scheduler Milestones | Epochs 5, 15, 30 |
| LR Decay Factor | 0.1 |
| Weight Decay | 1e-5 |
| Dropout Probability (FC layer) | 0.5 |
| Dropout Probability (Attention Layer) | 0.25 |
| Tile Size | 360 µm(1369 pixels) |

*Applied only during the training of ensemble models; the target model was trained without early stopping.

Supplementary Figure 1. A representative example of a slide in the Consistently Incorrect group (Clear Cell Carcinoma). Whole-slide image (top) with three regions highlighted at higher magnification (bottom). This slide contains only scattered tumor cells or small tumor clusters (boxed areas), lacking the subtype-defining histomorphological characteristic of CC. Such cases were therefore excluded from morphologic spectrum construction.

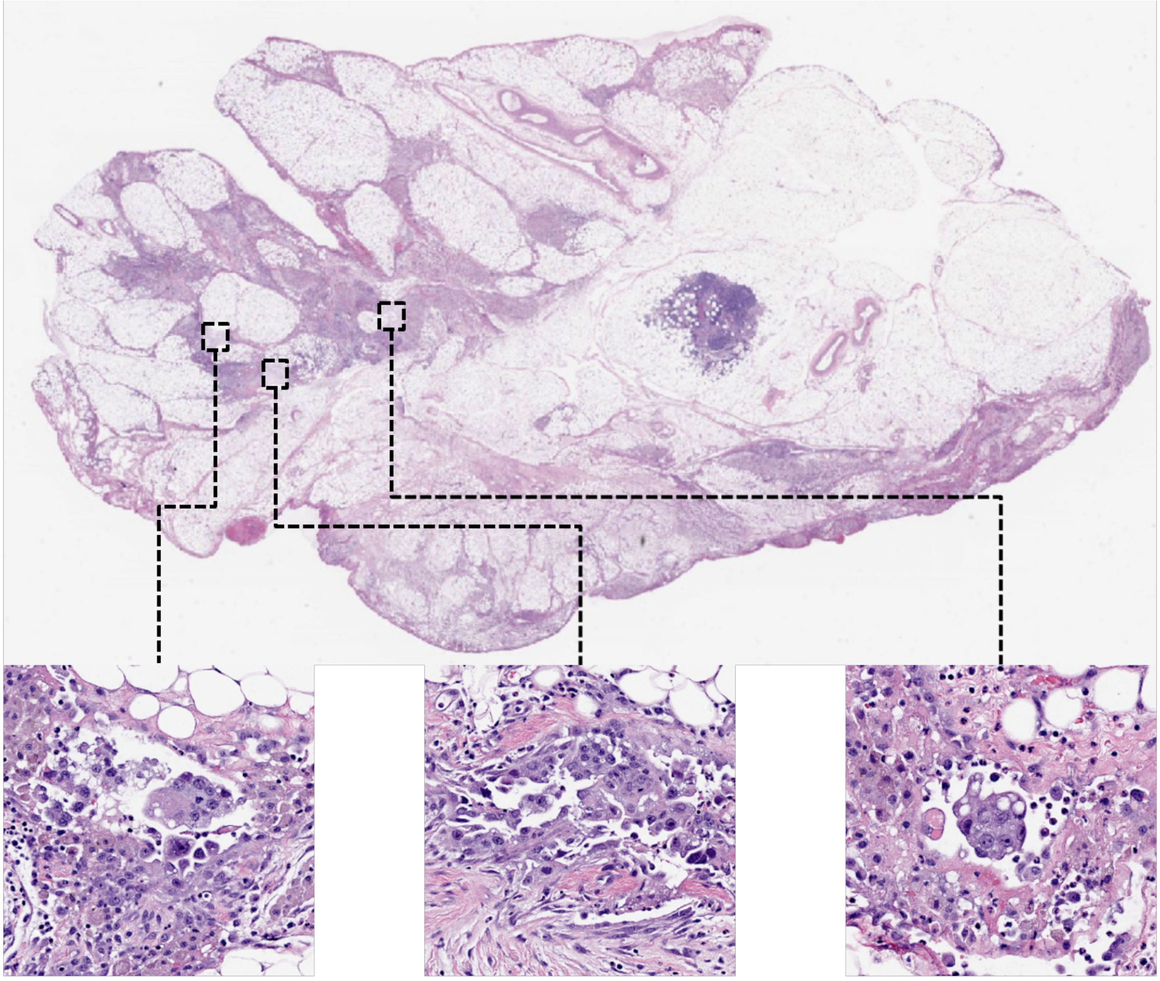

Supplementary Figure 2. Data partitioning and train/validation/test workflow. **a)** The ovarian tumor dataset was randomly shuffled at the patient level to generate five independent patient permutations. For each permutation, patients—together with all their slides—were distributed into ten partitions in a round-robin fashion. From these ten partitions, ten distinct 6:2:2 train/validation/test splits were created by cyclically designating two adjacent partitions as the test set. As a result, each partition appears in the test set twice within a permutation, so every slide is tested twice per permutation and ten times across the five permutations. **b)** For each data split, the model was trained on six training partitions and monitored on two validation partitions using an early-stopping strategy. The checkpoint with the lowest validation loss was used for inference on the two hold-out test partitions. **c)** Test results from all splits were collected, and performance metrics (e.g., AUC) were averaged across splits and permutations to obtain the final ensemble performance.

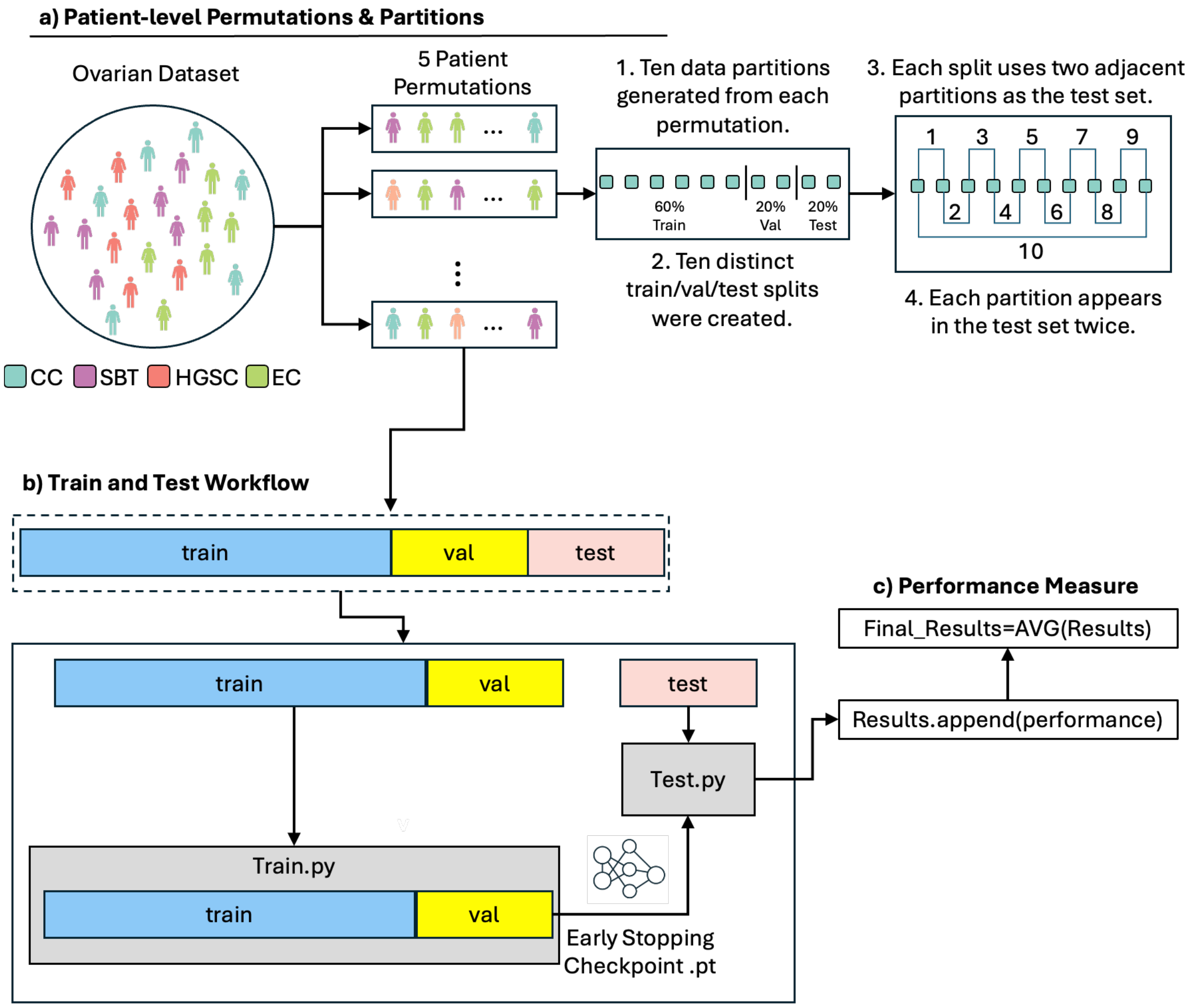

Supplementary Figure 3. Consensus clustering heatmaps for high-attention patches identified from Consistently Correct Slides in Clear Cell Carcinoma and Serous Borderline Tumor, across candidate cluster numbers (k = 2–8), quantified by Cophenetic Correlation Coefficient (COPH).

a) Clear Cell Carcinoma

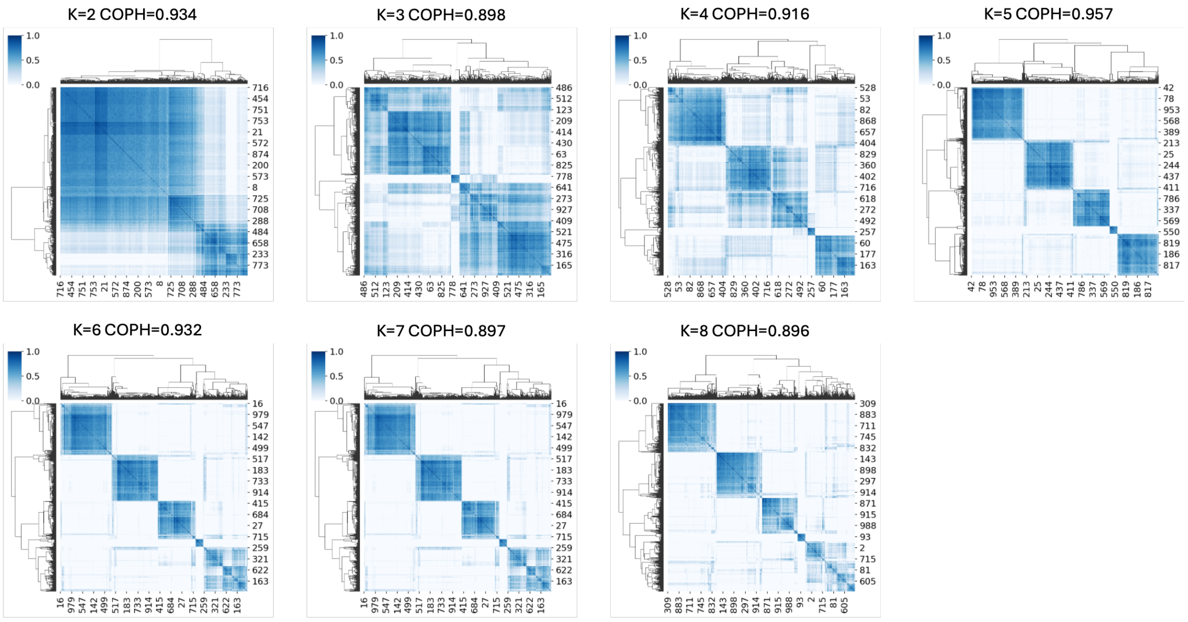


b) Serous Borderline Tumor

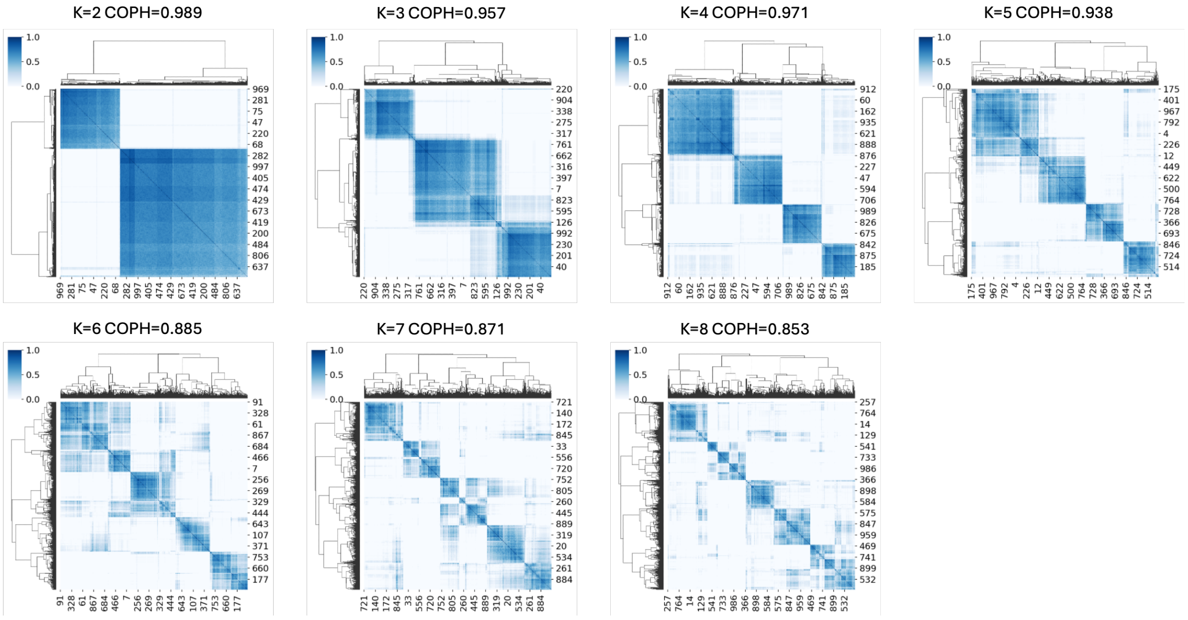

Supplementary Figure 4. Consensus clustering heatmaps for high-attention patches identified from Consistently Correct Slides in Endometrioid Carcinoma and High-grade Serous Carcinoma, across candidate cluster numbers (k = 2–8), quantified by Cophenetic Correlation Coefficient (COPH).

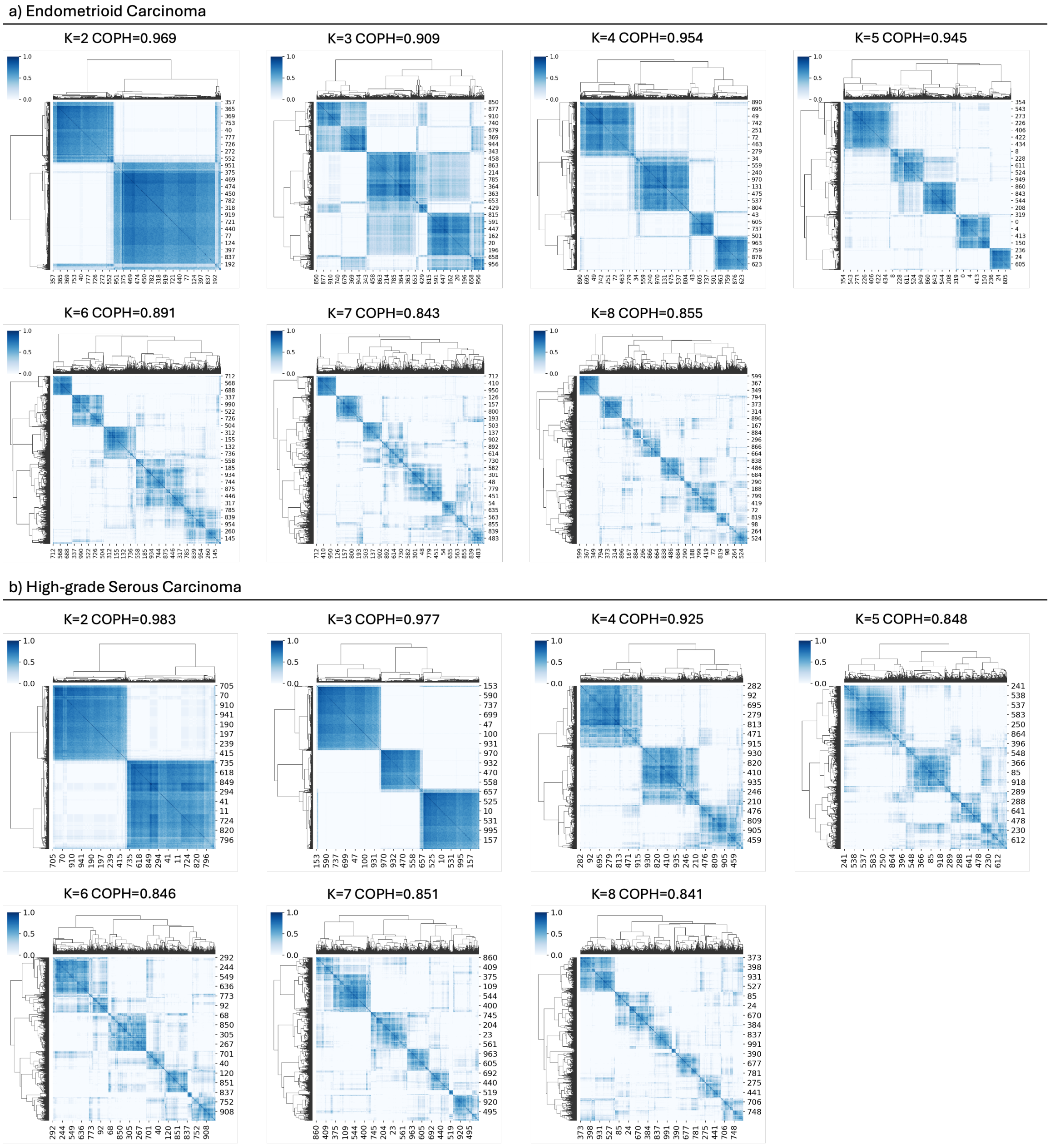

Supplementary Figure 5. Consensus clustering heatmaps for high-attention patches identified from Highly Variable slides, across candidate cluster numbers (k = 2–8), quantified by Cophenetic Correlation Coefficient (COPH).

a) Endometrioid Carcinoma → High-grade Serous Carcinoma

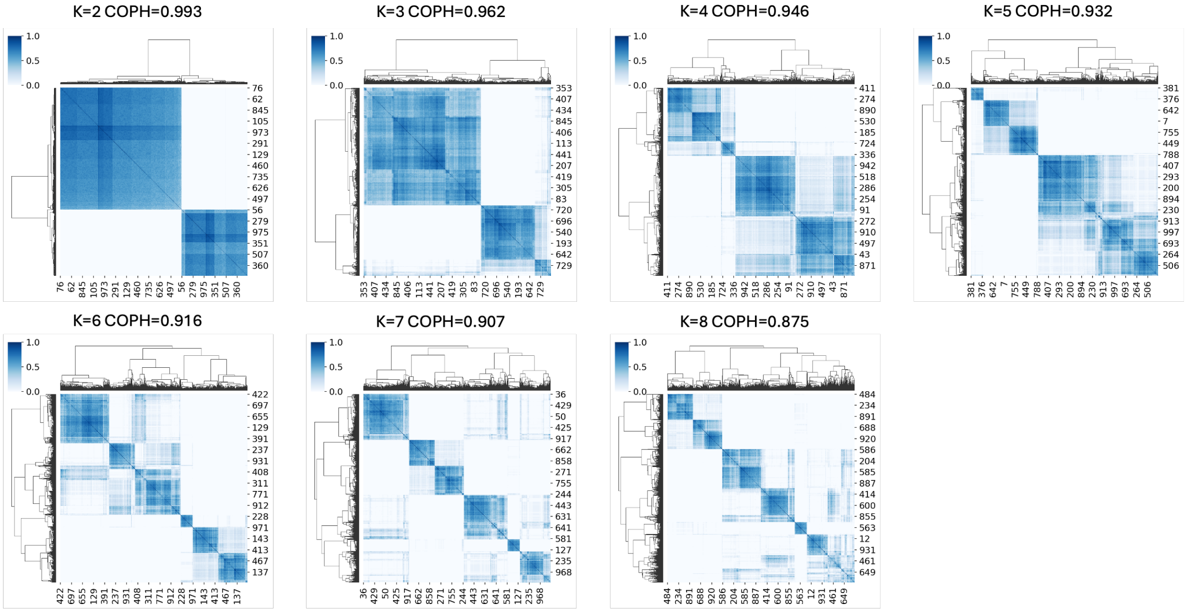


b) High-grade Serous Carcinoma → Endometrioid Carcinoma

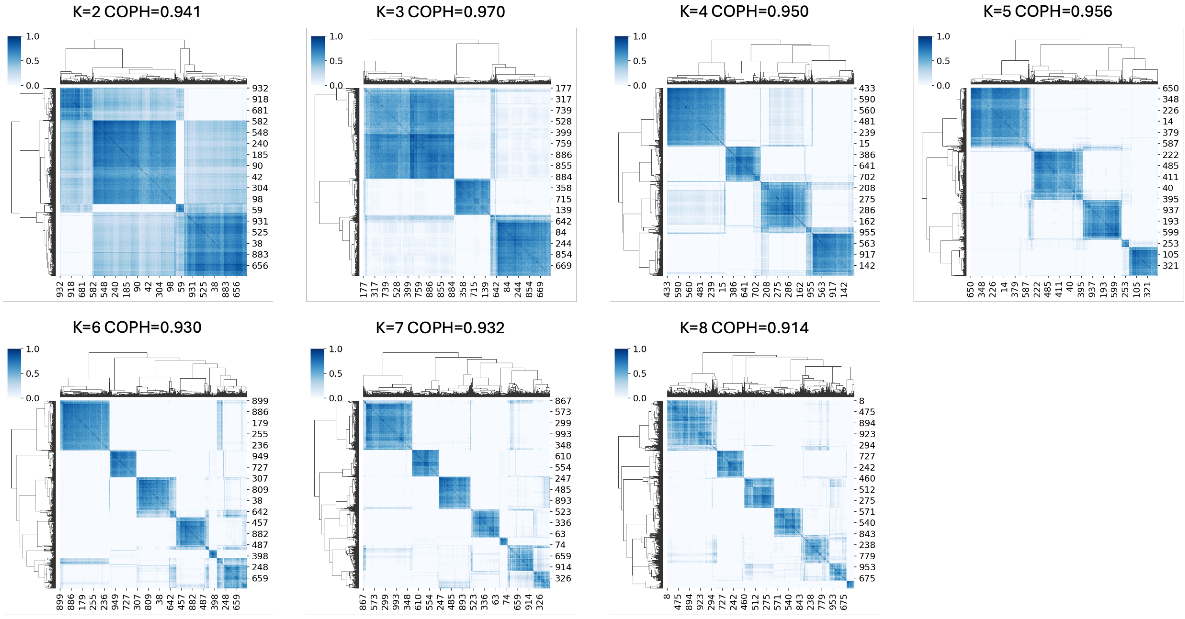

Supplementary Figure 6. Enrichment of High-Contribution Patches Across Individual Histomorphological Clusters

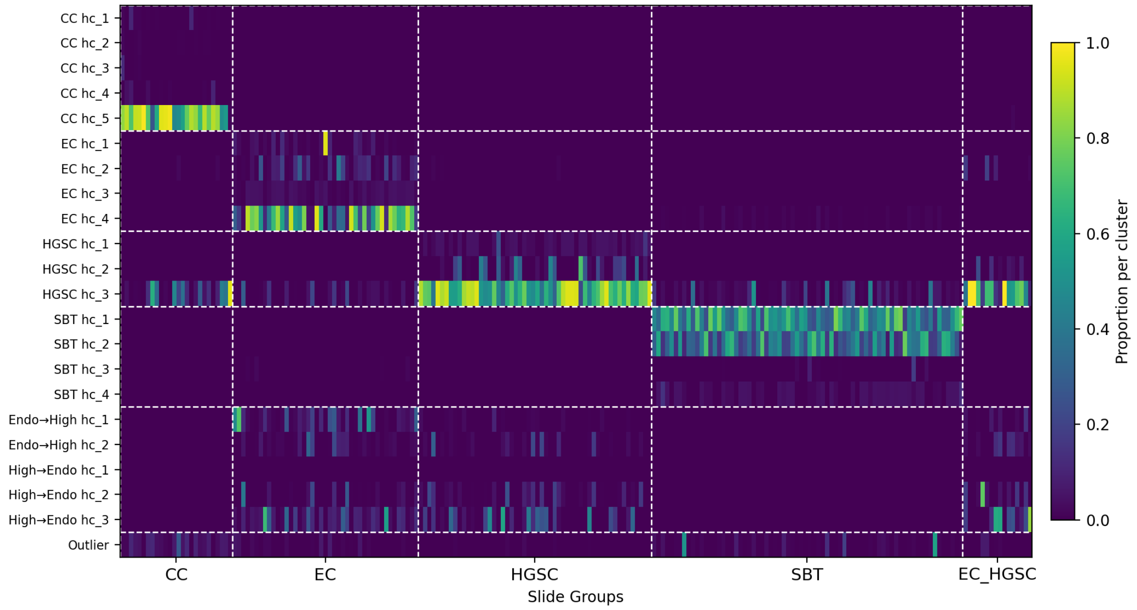

Supplementary Note 1: MorphoCA Manual

### 1.1 Install MorphoCA

To install MorphoCA, first launch QuPath, then drag the plugin file **qupath-extension-MorphoCA.jar** into the QuPath interface or install it via *Extensions* → *Install extension*. The JAR file can be downloaded from Github. After installation, QuPath needs to be restarted. Once restarted, the MorphoCA panel will be visible in the menu or sidebar (Figure 1).

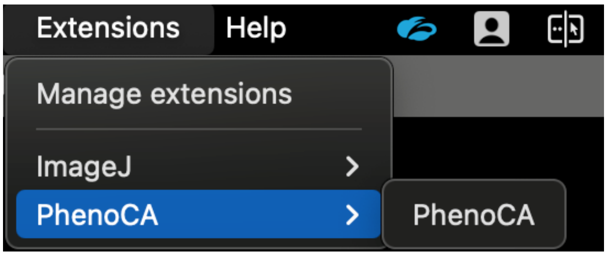


Figure 1 Installation of the MorphoCA extension in QuPath.

### 1.2 Load spatial coordinates

MorphoCA supports two modes of loading clustered patch coordinates: (i) single-slide import and (ii) project-level batch import, corresponding to the *Create File Selections* and *Create Project Selections* functions.

#### 1.2.1 Single-slide import (Create File Selections)

In the single-slide mode, users need to open the target slide in QuPath and load a CSV file[1] that contains the spatial coordinates of clustered patches. This file records the locations of high-contribution patches within the slide. The CSV typically includes the following fields: slide_name (ending with .svs and matching the filename of the opened slide), patch_id (identifier of each patch, e.g., patch_2316), hc_label (cluster label, such as hc_1, hc_2, etc.), x and y (top-left coordinates of the patch), as well as width and height (the width and height of the patch). An example is shown in Figure 2.

| A | B | C | D | E | F | G |
|---|---|---|---|---|---|---|
| ANON0LN0DJ16A_1_1.svs | patch_2316 | hc_1 | 100679 | 14940 | 1367 | 1367 |
| ANON0LN0DJ16A_1_1.svs | patch_2224 | hc_1 | 97944 | 9470 | 1367 | 1367 |
| ANON0LN0DJ16A_1_1.svs | patch_2235 | hc_1 | 99311 | 16307 | 1367 | 1367 |
| ANON0LN0DJ16A_1_1.svs | patch_2313 | hc_1 | 100679 | 13572 | 1367 | 1367 |
| ANON0LN0DJ16A_1_1.svs | patch_2234 | hc_1 | 99311 | 14940 | 1367 | 1367 |
| ANON0LN0DJ16A_1_1.svs | patch_2312 | hc_1 | 100679 | 12205 | 1367 | 1367 |
| ANON0LN0DJ16A_1_1.svs | patch_2401 | hc_1 | 103414 | 16307 | 1367 | 1367 |
| ANON0LN0DJ16A_1_1.svs | patch_2403 | hc_1 | 104781 | 16307 | 1367 | 1367 |
| ANON0LN0DJ16A_1_1.svs | patch_2227 | hc_1 | 99311 | 10837 | 1367 | 1367 |
| ANON0LN0DJ16A_1_1.svs | patch_2232 | hc_1 | 97944 | 14940 | 1367 | 1367 |
| ANON0LN0DJ16A_1_1.svs | patch_2193 | hc_1 | 95209 | 49127 | 1367 | 1367 |
| ANON0LN0DJ16A_1_1.svs | patch_1799 | hc_1 | 85636 | 10837 | 1367 | 1367 |
| ANON0LN0DJ16A_1_1.svs | patch_2442 | hc_1 | 104781 | 42290 | 1367 | 1367 |
| ANON0LN0DJ16A_1_1.svs | patch_2437 | hc_1 | 103414 | 40922 | 1367 | 1367 |

Figure 2 Example of a CSV file containing spatial coordinates and cluster labels of patches.

After selecting the CSV file with *SelectFile* in the MorphoCA panel, the 'In File:' field displays the imported coordinate filename, whereas the 'Out File:' field specifies the filename for storing annotation results (see the following section "Saving Annotations"). By then clicking *Create File Selections*, the extension overlays the patches from the CSV onto the current slide as colored boxes or regions. Different clusters (hc_label) are displayed in distinct colors (Figure 3).

[1] For details on how to prepare the CSV file, please refer to the instructions provided in the GitHub README.

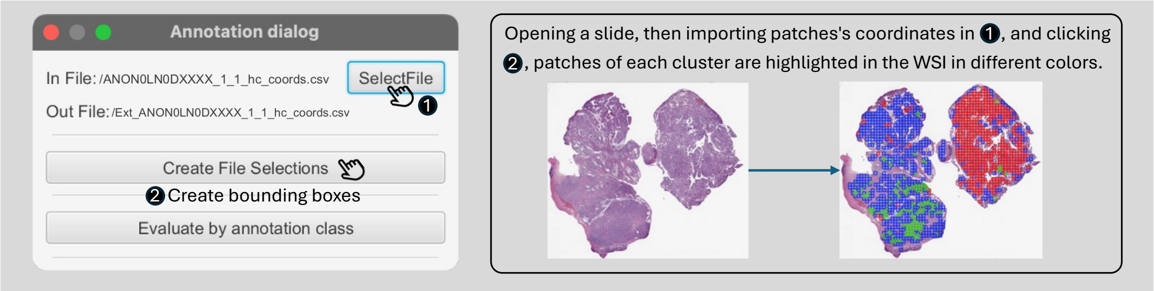


Figure 3 Example of loading a CSV file and visualizing clustered patches with distinct colors.

By default, QuPath also visualizes the patch_id on each patch after the coordinates are loaded (Figure 4), which may result in visual clutter. If the display of patch_id is not needed, users can disable it by clicking *View → Show annotation names*.

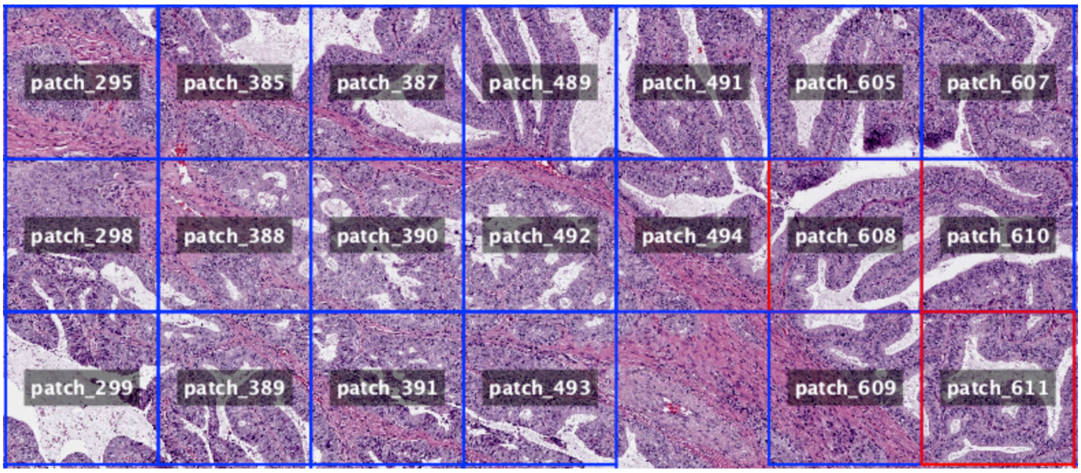


Figure 4 Visualization of clustered patches in QuPath, with patch IDs displayed by default.

1.2.2 Create Project Selections

In large-scale data scenarios, MorphoCA supports batch visualization by creating a QuPath project (Figure 5). The procedure involves merging the spatial coordinate CSV files of clustered patches from multiple slides into a single file. This file must contain a *slide_id* field that exactly matches the slide names in the QuPath project, while the other columns follow the same format as in the single-slide mode. Once a new project is created in QuPath and all relevant slides are imported, users can select the merged CSV file in the MorphoCA panel and click *Create Project Selections*. The extension will then automatically distribute the coordinates to each slide according to the *slide_id* and overlay the clustered results in distinct colors on the corresponding WSIs. This functionality is particularly useful for large cohorts, as it supports efficient batch visualization and streamlined annotation.

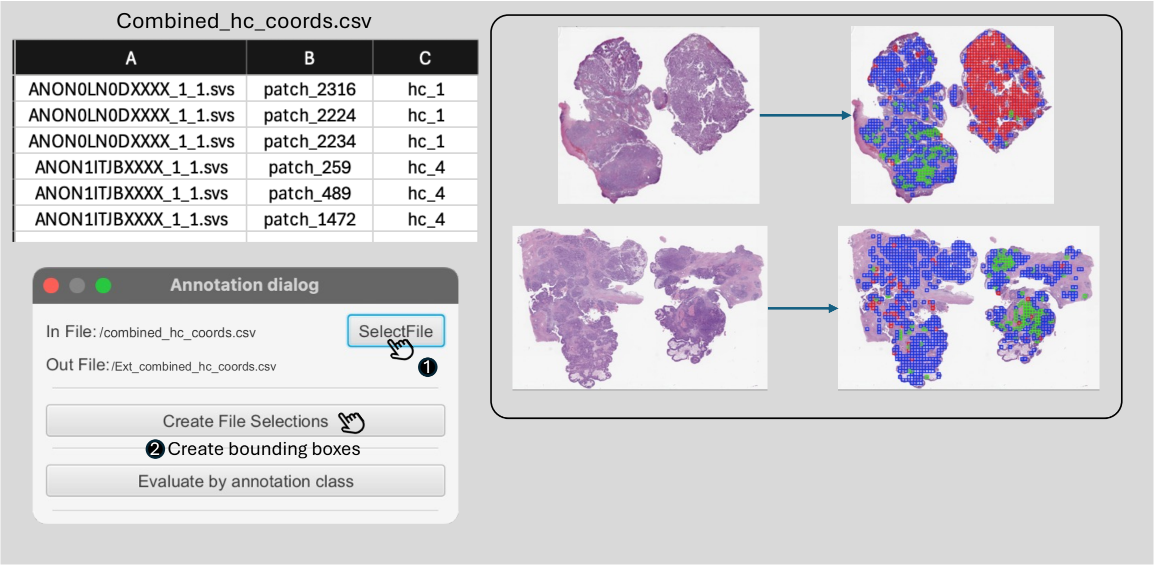


Figure 5 Example of batch visualization of clustered patches in MorphoCA.

### 1.3 Interactive Annotation

#### 1.3.1 Entering Annotation Mode

After the spatial coordinates have been loaded in MorphoCA, users can click *Evaluate by annotation class* to enter the annotation mode (Figure 6). By selecting a cluster from the class list (for example, hc_1) and then clicking *Process selected class*, the chosen cluster will remain highlighted while other clusters are automatically hidden. This allows users to focus on the distribution and morphological features of the selected cluster patches. At this stage, users enter a whole-slide contextual environment for visualization and annotation, where they can freely zoom or pan to examine features at both the cellular and tissue levels.

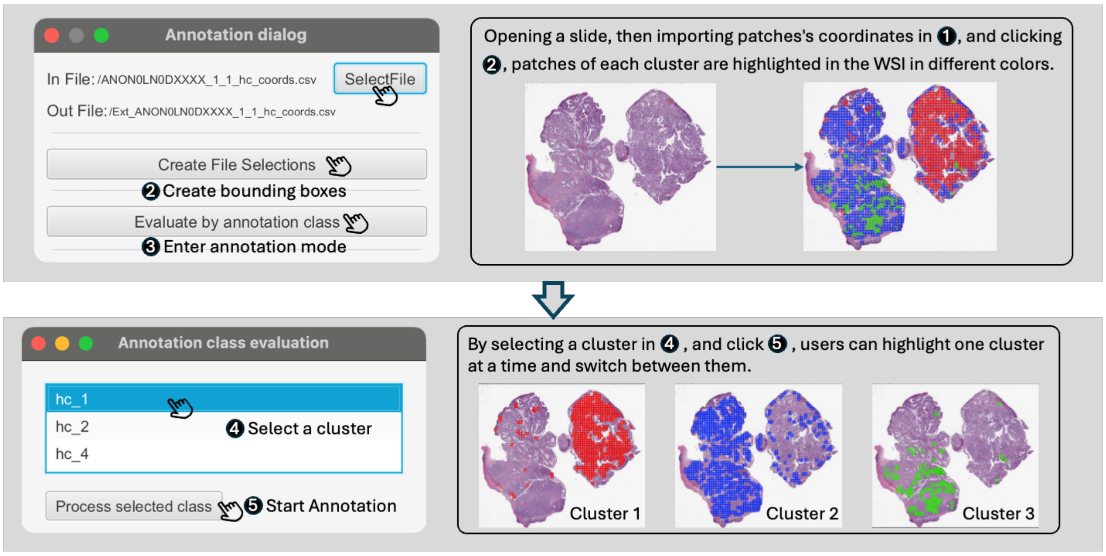


Figure 6 Entering annotation mode in MorphoCA

#### 1.3.2 Multiscale Review

At high magnification, users can examine the cytological features of individual patches. At low magnification, they can evaluate the spatial distribution of the clustered patches across the whole slide and their relationship with histological structures. This combination of high-

and low-power views provides a comprehensive morphological context and forms the basis for subsequent annotation questions (Figure 7 right).

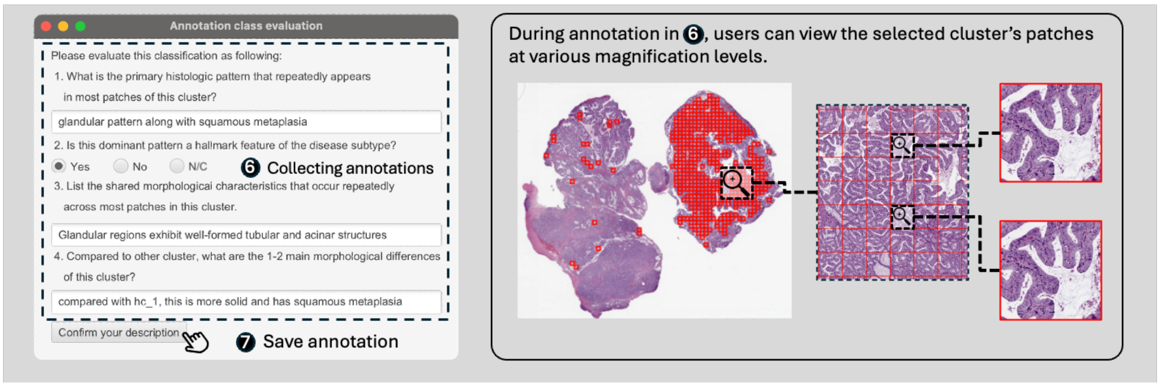


Figure 7 Workflow of annotation in MorphoCA

#### 1.3.3 Four Annotation Questions

In the pop-up panel, users are required to fill in four types of information for the current cluster (Figure 7 left). The first is **Represented Phenotype**, which describes the main phenotype of the cluster, recorded using standard terminology (e.g., glandular architecture, papillary architecture, solid pattern, squamous metaplasia). If uncertain, users should input **N/C** (Not Certain). The second is **Subtype-Defining Phenotype**, which determines whether the phenotype is a core diagnostic feature for the subtype. The options are Yes, No, or N/C, and the judgment should follow diagnostic criteria or guidelines. The third is **Detailed Morphological Descriptions (Intra-cluster Features)**, which require listing the consistent detailed features that appear across most patches in the cluster, such as prominent nucleoli, pleomorphism, or stromal fibrosis. The last is **Inter-cluster Morphological Differences**, which compare the current cluster with other clusters of the same phenotype, requiring 2–3 distinguishing traits. For example, "compared with other glandular clusters, this cluster shows richer stroma or denser solid regions." If uncertain, users should enter N/C. It is important to note that responses should reflect features consistently observed across the majority of patches, rather than being biased by a few outliers..

#### 1.3.4 Saving Annotations

Once the four questions have been completed, users can click *Confirm your description* to save the structured annotation results locally. A new CSV file will be created in the same directory as the imported CSV, with filenames beginning with 'Ext_'. Users should repeat the same process for all clusters and slides. In batch annotation scenarios, it is recommended to save after completing several clusters at a time to reduce the risk of data loss caused by accidental operations or software crashes.

### Supplementary Note 2: MorphoExplainer Manual

#### 2.1 Install MorphoExplainer

To install MorphoExplainer, first launch QuPath, then drag the plugin file **qupath-extension-MorphoExplainer.jar** into the QuPath interface or install it via *Extensions → Install extension.* The JAR file can be downloaded from Github. After installation, QuPath needs to be restarted. Once restarted, the MorphoExplainer panel will be visible in the menu or sidebar.

#### 2.2 Loading GeoJSON Files

MorphoExplainer reads **GeoJSON files** that contain overlay information for both attention heatmaps and spectrum-based explanations. Each GeoJSON file corresponds to a single WSI.

##### 2.2.1 GeoJSON File Schema

Each WSI is saved as one GeoJSON FeatureCollection that contains three logical parts, as illustrated in Figure 8.

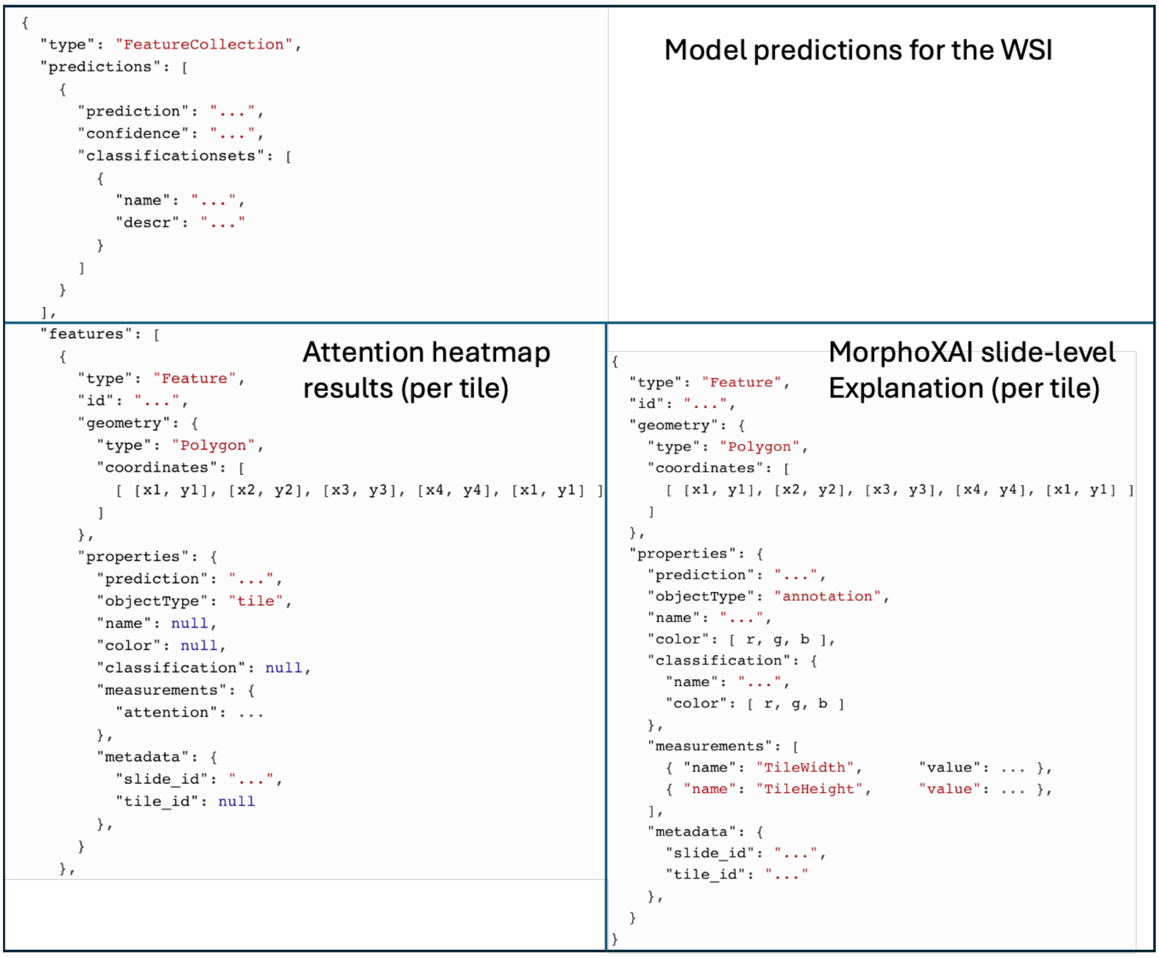


Figure 8 GeoJSON file schema used by MorphoExplainer

The first part is Model predictions for the WSI, which records the model's four-class prediction outputs for the given slide. Because the model used in this study performs a four-class classification task, this array contains four elements, each corresponding to one disease subtype. For each element, the attribute **prediction** specifies the subtype label, **confidence** stores the model's predicted probability for that subtype, and **classificationsets** records the morphologic clusters that the model's high-contribution patches were mapped to for that subtype. Within each entry of **classificationsets**, **name** denotes the cluster identifier (for example, Endo_hc_2), and **descr** provides a concise textual description of the cluster's morphologic characteristics (for example, Glandular pattern with squamous metaplasia). The cluster identifiers listed here are referenced by the morphologic spectrum features described later.

The second part, Attention heatmap results (per tile), stores tile-level information for the model's attention visualization. Each element within the **features** array represents one tile from the WSI. The spatial boundary of the tile is defined in the **geometry** field as a polygon described by its pixel coordinates. Within the **properties** field, **prediction** indicates which of the four predicted classes this tile belongs to, and **objectType** is set to "tile." The **measurements** subfield contains the tile's attention weight (**attention**), which quantifies the relative contribution of that tile to the model's prediction. The **metadata** field records the associated slide identifier (**slide_id**) and, when available, the tile identifier (**tile_id**).

The third part, Morphologic spectrum interpretability results (per tile), also appears under the **features** array but represents the model's spectrum-based explanations. Each feature again corresponds to an individual tile, with its spatial boundary defined in the same **geometry** format as above. In the **properties** field, **prediction** specifies the subtype to which this tile contributes, and **objectType** is set to "annotation." The **name** attribute identifies the cluster element, while **color** defines the RGB color used to display the tile on the WSI. The **classification** subfield stores the tile's morphologic cluster assignment, where **name** corresponds to the cluster identifier that matches one of the entries in the top-level **classificationsets**, and **color** provides the display color for that cluster. The **measurements** array contains quantitative information such as tile width, tile height. Finally, the **metadata** field again records the slide and tile identifiers.

The GeoJSON files containing these interpretability results for each WSI can be automatically generated using the code provided with this study; implementation details and scripts are available on the project's GitHub repository.

##### 2.2.2 Import GeoJSON into MorphoExplainer

To import a GeoJSON file: Open the corresponding WSI in QuPath, launch MorphoExplainer, and click the Select GeoJSON File button. In the file selection window, choose the GeoJSON file you wish to import. Once selected, the file will be automatically loaded into the plugin and its contents displayed in the corresponding visualization panels.

#### 2.3 Interactive Annotation for Visualization and Rating

MorphoExplainer provides three interactive panels—**Heatmap**, **MorphoSpectrum**, and **MorphoSpectrum Cluster**—that allow users to explore the model's attention heatmaps and morphologic spectrum–based explanations, as well as to provide corresponding annotations and ratings.

##### 2.3.1 Heatmap Panel

Clicking the **Heatmap** tab enters the Heatmap Panel (Figure 9), which visualizes the model's attention heatmap on the whole-slide image. To begin, users should click the **Attention Heatmap Select** button, prompting the plugin to read the attention-related information stored in the imported GeoJSON file and overlay the corresponding heatmap on the WSI.

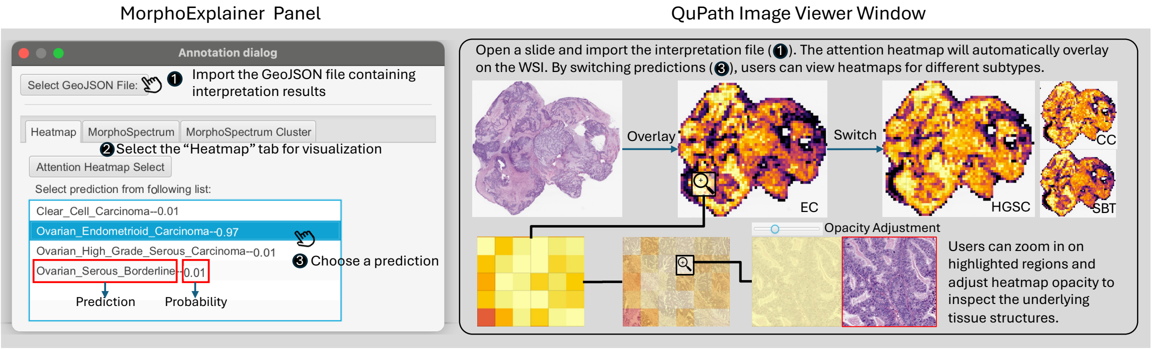


Figure 9 Interface and workflow of the Heatmap panel in MorphoExplainer

In the blue box below the panel, the plugin lists all possible predictions together with their associated predicted probabilities. Selecting one prediction will automatically update the QuPath viewer to display the attention heatmap corresponding to that subtype. The heatmap highlights the regions that contributed most strongly to the model's decision. Using QuPath's opacity slider in the viewer toolbar, users can adjust the transparency of the heatmap layer to better visualize the underlying histologic structures. Users can also zoom in on the highlighted areas to closely inspect tissue morphology corresponding to high-attention regions.

At the bottom of the panel (Figure 10), three structured questions are presented for the annotator to evaluate the interpretability and diagnostic usefulness of the attention heatmap. Annotators can select their responses corresponding to each question. After completing the ratings, clicking **"Confirm your description"** will automatically save the annotation results.

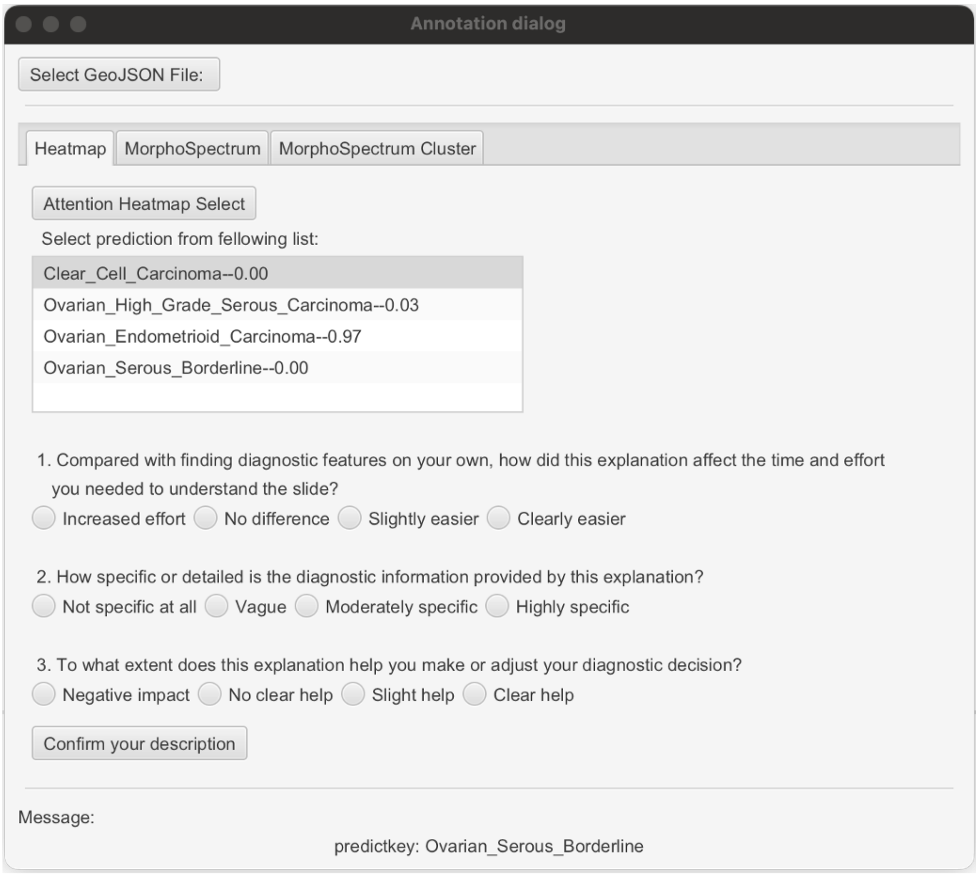


Figure 10 Annotation interface for the Heatmap panel in MorphoExplainer

#### 2.3.2 MorphoSpectrum Panel

Clicking the **MorphoSpectrum** tab opens the MorphoSpectrum panel (Figure 11), which visualizes the model's spectrum-based interpretability results on the WSI. After clicking the **MorphoSpectrum Select** button, the plugin reads the spectrum-related information from the imported GeoJSON file and overlays the morphologic spectrum interpretation on the slide.

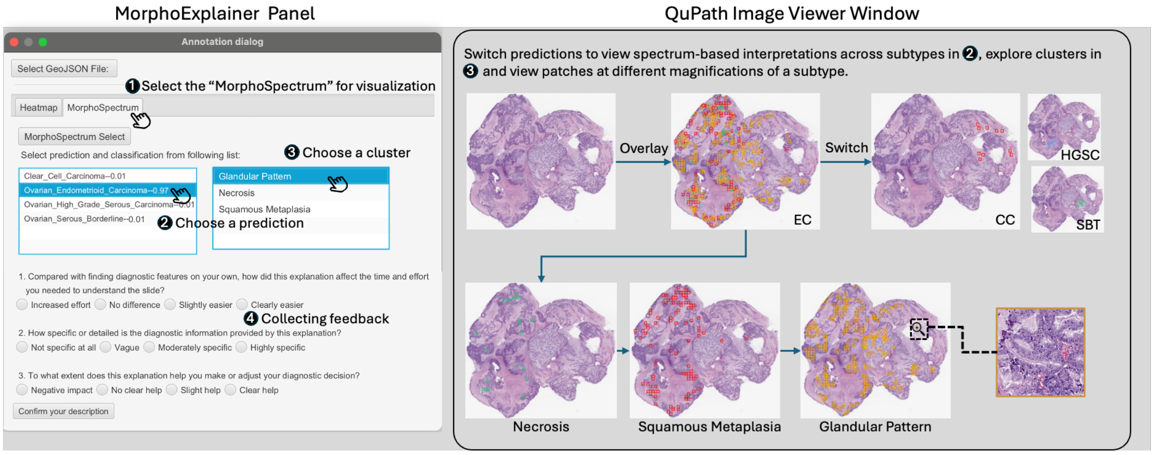


Figure 11 Interface and workflow of the MorphoSpectrum panel in MorphoExplainer

In the lower section of the panel, two blue selection boxes are provided. The left box lists all subtype predictions made by the model; users can switch between them to view the spectrum-based explanation corresponding to each subtype. The right box lists the morphologic clusters that constitute the selected subtype's spectrum, each labeled with its morphologic description (for example, Glandular pattern, Squamous metaplasia, or Necrosis). Clicking on a cluster highlights its corresponding patches on the WSI, allowing users to inspect where and how these morphologic patterns are distributed across the tissue.

At the bottom of the panel, three structured questions are provided for annotators. After completing the ratings, clicking **"Confirm your description"** will automatically save the annotation results.

#### 2.3.3 MorphoSpectrum Cluster Panel

The MorphoSpectrum Cluster panel provides the same visualization functions as the MorphoSpectrum panel. The only difference lies in the annotation component. This panel is designed specifically for pathologists to evaluate whether the highlighted patches visually correspond to the described morphologic pattern of each cluster. At the bottom of the panel, a single structured question is provided—"Do the highlighted patches correspond to the described morphological patterns?"—with three selectable options: Consistent, Inconsistent, and Uncertain. After choosing the appropriate option, users can click "Confirm your description" to save their evaluation for the current cluster.

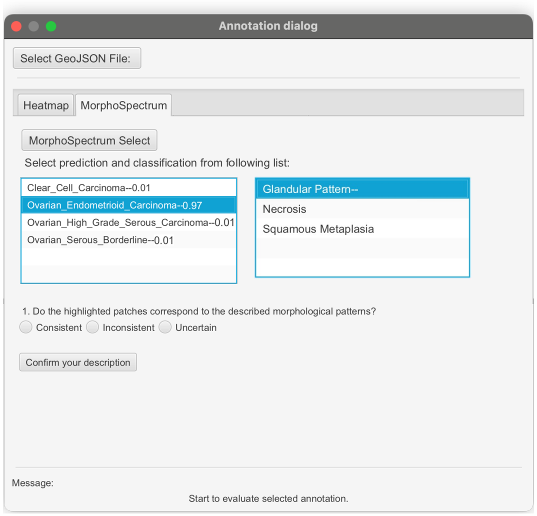


Figure 12 Interface of the MorphoSpectrum Cluster panel

### 2.4 Annotation Results

For each WSI, the annotations provided for both the attention heatmap and the morphologic spectrum explanations are automatically exported as a .csv file (Figure 13), which is saved in the same directory as the opened .svs file. In the exported file, the first column records the slide filename. The first two rows correspond to the annotator's ratings of the attention heatmap and the morphologic spectrum explanation regarding ease of understanding, specificity of diagnostic information, and helpfulness for diagnostic decision-making. The subsequent rows contain the evaluations for each morphologic cluster, indicating whether the highlighted patches are morphologically consistent with the described cluster pattern.

| A | B | C | D | E | F |
|---|---|---|---|---|---|
| slide.svs | Heatmap | | Clearly easier | Moderately specific | Slight help |
| slide.svs | MorphoSpectrum | | Clearly easier | Highly specific | Clear help |
| slide.svs | MorphoSpectrum Cluster | Endo_hc_1--Squamous Metaplasia | Consistent | | |
| slide.svs | MorphoSpectrum Cluster | Endo_hc_2--Glandular Pattern with Squamous Metaplasia | Consistent | | |
| slide.svs | MorphoSpectrum Cluster | Endo_hc_4--Glandular Pattern | Consistent | | |
| slide.svs | MorphoSpectrum Cluster | High_to_Endo_hc_3--Glandular Pattern | | | |

Figure 13 Interface of the MorphoSpectrum Cluster panel

Supplementary Note 3: Pathologist Questionnaire for Interpretability Evaluation

Q1. Ease of Understanding (Cognitive Effort)

Compared with finding diagnostic features on your own, how did this explanation affect the time and effort you needed to understand the slide?

| Score | Description |
|---|---|
| 0 - Increased effort | It required more time and effort to understand than interpreting the slide on my own. |
| 1 - No difference | The time and effort needed were about the same as interpreting on my own. |
| 2 - Slightly easier | It helped me understand a bit faster or with less effort. |
| 3 – Clearly easier | It made understanding noticeably faster and easier than interpreting on my own. |

Q2. Specificity of Diagnostic Information (Diagnostic Informativeness)

How specific or detailed is the diagnostic information provided by this explanation?

| Score | Description |
|---|---|
| 0 - Not specific at all | The explanation shows no meaningful diagnostic information. |
| 1 - Vague | It roughly points out areas of interest, but their diagnostic meaning is unclear. Not sure what phenotypes are used for diagnosis. |
| 2 - Moderately specific | The diagnostic features are intuitively recognizable but not clearly defined. |
| 3 – Highly specific | The diagnostic features are well recognizable and clearly defined. |

Q3. Impact on Diagnostic Decision (Actionability)

To what extent does this explanation help you make or adjust your diagnostic decision?

| Score | Description |
|---|---|
| 0 - Negative impact | It misleads or distracts me; I would rather make the judgment without this explanation. |
| 1 - No clear help | The explanation does not help me make or adjust my diagnostic judgment. |
| 2 - Slight help | It provides some guidance, but not sufficient for confident decision making. |
| 3 – Clear help | It clearly directs me to meaningful diagnostic features—either confirming the diagnosis or revealing atypical findings that make me reconsider it. |
